# Spinal cerebrospinal fluid contacting neurons are a conserved sensory neuronal population along the mouse spinal cord

**DOI:** 10.1101/2024.10.10.617653

**Authors:** Crozat Elysa, Blasco Edith, Ramirez-Franco Jorge, Riondel Priscille, Jurčić Nina, Seddik Riad, Michelle Caroline, Trouslard Jérôme, Wanaverbecq Nicolas

**Author notes:** Correspondence should be addressed to (WN). CE, BE & RFJ Contributed equally to this work as first authors.

## Abstract

Cerebrospinal Fluid contacting neurons (CSF-cNs) are present in all vertebrates around the medullo-spinal central canal. They are GABAergic, selectively express PKD2L1, a member of the TRP channel superfamily, and are thought to represent a novel sensory system intrinsic to the central nervous system. Using histology, we found that CSF-cNs form a homogeneous population, distributed along the whole spinal cord and mainly located in the ventral region of the central canal. Patch-clamp recordings reveal conserved intrinsic properties and voltage-dependent conductance expression. Spinal CSF-cNs express PKD2L1 channels and ASICs, acting as sensory receptors for extracellular pH changes. They express both inhibitory (GABA_A_, glycine) and excitatory (glutamate, cholinergic) synaptic receptors as well as functional GABA_B_ and muscarinic receptors, but not glutamatergic metabotropic ones, to modulate Ca^2+^ channels.

CSF-cNs represent a functionally homogeneous population that might integrates sensory signals along the central canal to modulate body functions by regulating local spinal networks. Demonstrating such a function represents the future challenge in the field.

## INTRODUCTION

Neurons in contact with cerebrospinal fluid (CSF-cNs) are present around the central canal (CC) in numerous vertebrate species^1–9^ but remain functionally under-characterized^10^ in mammals. These GABAergic bipolar neurons have a unique morphology, with their soma located beneath or within the ependymocyte monolayer lining the CC^1–9^. They project a single dendrite toward the CC lumen, ending in a ciliated protrusion that contacts the CSF^7,9,11,12^. Their axon extends into the ventral spinal cord (SC), and form long, bilateral fiber bundles at the median fissure^5,11,13–15^. Unlike the long range sparse GABAergic projections^16^ observed in the hippocampus^17^ and the inferior olive^18^, CSF-cNs form a unique dense network of ascending fibers running along several SC segments and extending collaterals within the spinal tissue with recurrent synapses onto CSF-cNs^14,15^.

In juvenile rats^9^, CSF-cNs exhibit spontaneous action potential (AP) firing, mediated by sodium and potassium voltage-dependent channels, and they also express voltage-dependent calcium channels (Ca_V_ 2.2 and Ca_V_ 3.1-3)^19,20,21^. CSF-cNs appear integrated in a neuronal network receiving primarily inhibitory inputs (GABA_A_ and glycine receptors)^12,19^, as well as excitatory ones (AMPA/kainate receptors)^6,19^. They also exhibit homo- and heterosynaptic modulations through metabotropic GABAergic receptors (GABA_B_-Rs)^19^. These neurons express PKD2L1, a channel sensitive to pH, and osmolarity changes (chemosensitivity)^12,22–25^ as well as CSF flow, and SC bending (mechanosensitivity)^26–28^. Additionally, they express acid-sensing ion channels (ASICs) and both channels are capable of modulating CSF-cN excitability^22,24^

Morphological^6,8,29,30^ and phenotypic^8,30–32^ differences suggest CSF-cNs are regrouped in two subpopulations. CSF-cNs present in the CC ventral region exhibit immature phenotypes^5,8,29–31^ combined with single AP firing^8,30^. Dorsolateral CSF-cNs have a more mature phenotype and tonic AP discharge^8,29–31^. In lumbar segments, the two subpopulations also differ in their responses to GABAergic signaling, either inducing depolarization or hyperpolarization^33^.

In zebrafish larvae, CSF-cNs selectively activate motor neurons and interneurons, influencing swimming behavior^26,34,35^. In mice, they play a role in controlling posture, balance^15^, and locomotion^14^. Finally, they have been shown to respond to bacterial toxins, suggesting a potential role in the body’s immune defense^36^. More recently, it was shown that, following SC injury, CSF-cN constitutive activation through κ-opioid signaling is halted leading to disinhibition of ependymal cell proliferation to promote scar formation^10^.

Anatomical and functional evidence suggest that CSF-cNs serve as a novel sensory or interoceptive system intrinsic to the CNS. Due to their distinct properties and distribution along the SC, these neurons may integrate into specific spinal networks to modulate body functions.

This study investigated, in the mouse SC, the distribution, morphology, and properties of CSF-cNs, a neuronal population that still remains enigmatic. Histological analyses reveal that CSF-cNs form a dense population along the SC axis, primarily located in the CC ventral region. Electrophysiological recordings show that they share similar intrinsic properties along the SC and express functional GABA_A_, glycine, glutamate and cholinergic receptors. Additionally, functional GABA_B_ and muscarinic receptors, but not glutamatergic metabotropic ones, modulate Ca^2+^ channels activity.

***CSF-cNs would represent a functionally homogenous neuronal population integrating sensory signals to modulate specific spinal networks and body functions both in physiological and pathological conditions. Further research is required to characterize in-depth CSF-cN functional connectivity combined with behavioral studies. This represents the future challenge in the field to gain a deeper understanding of their integration into local spinal networks and function(s)***.

## RESULTS

### CSF-cN morphology and distribution along the central canal axis

CSF-contacting neurons (CSF-cNs) are found along the entire CC in various vertebrate species, exhibiting a consistent morphology^1–9^. In this study, we assessed the distribution and density of CSF-cNs in the mouse SC using the PKD2L1-Cre::tdTomato mouse model, to selectively label CSF-cNs with fluorescent tdTomato protein. CNS tissue was cleared using the vDISCO^37,38^ method and imaged *via* light sheet microscopy, allowing 3D visualization of the entire SC. ***Figure 1A*** shows the anatomy of the CNS (tissue autofluorescence) and the localization of CSF-cNs along the CC (red line). Imaging of cervical, thoracic, and lumbar regions at higher magnification (Boxes 1-3 in ***Fig. 1A, Bottom***) reveals CSF-cN cell bodies distributed along the CC in all segments (***Fig. 1B-D***). Their axons projected into the ventral SC, forming bilateral fiber bundles ***(Fig*.*1B-D, Right*** ventral view). These neurons were also found in distal ventral regions, consistent with previous studies^13^. 3D reconstruction images were segmented to quantify CSF-cN cell bodies (yellow objects) as well as their axonal projection (labelled in blue; ***Fig*.*1B-D, Bottom***). To better resolve CSF-cN distribution and quantification along the CC axis, we acquired images using a higher magnification objective (12x; see ***Supplementary Figure 1***). Due to the small size of CSF-cNs, clustering tendencies, and tissue shrinkage from the clearing method, we faced technical limitations and reached microscopy optical limits to resolve and identify single CSF-cN somatas (see ***Supplementary Figure 2***). Further analysis was carried out using thin sections and confocal microscopy (***Figure 2A***). Results show that CSF-cNs are present in all SC segments but at a higher density in the lumbar compared to thoracic and cervical regions (***Fig. 2B***). A larger proportion of CSF-cNs is located in the ventral CC (***Fig. 2A, Right; 2C***) and neurons are closer to the CC in cervical and lumbar segments but further away in the thoracic one (***Fig. 2D***).

**Figure 1.**
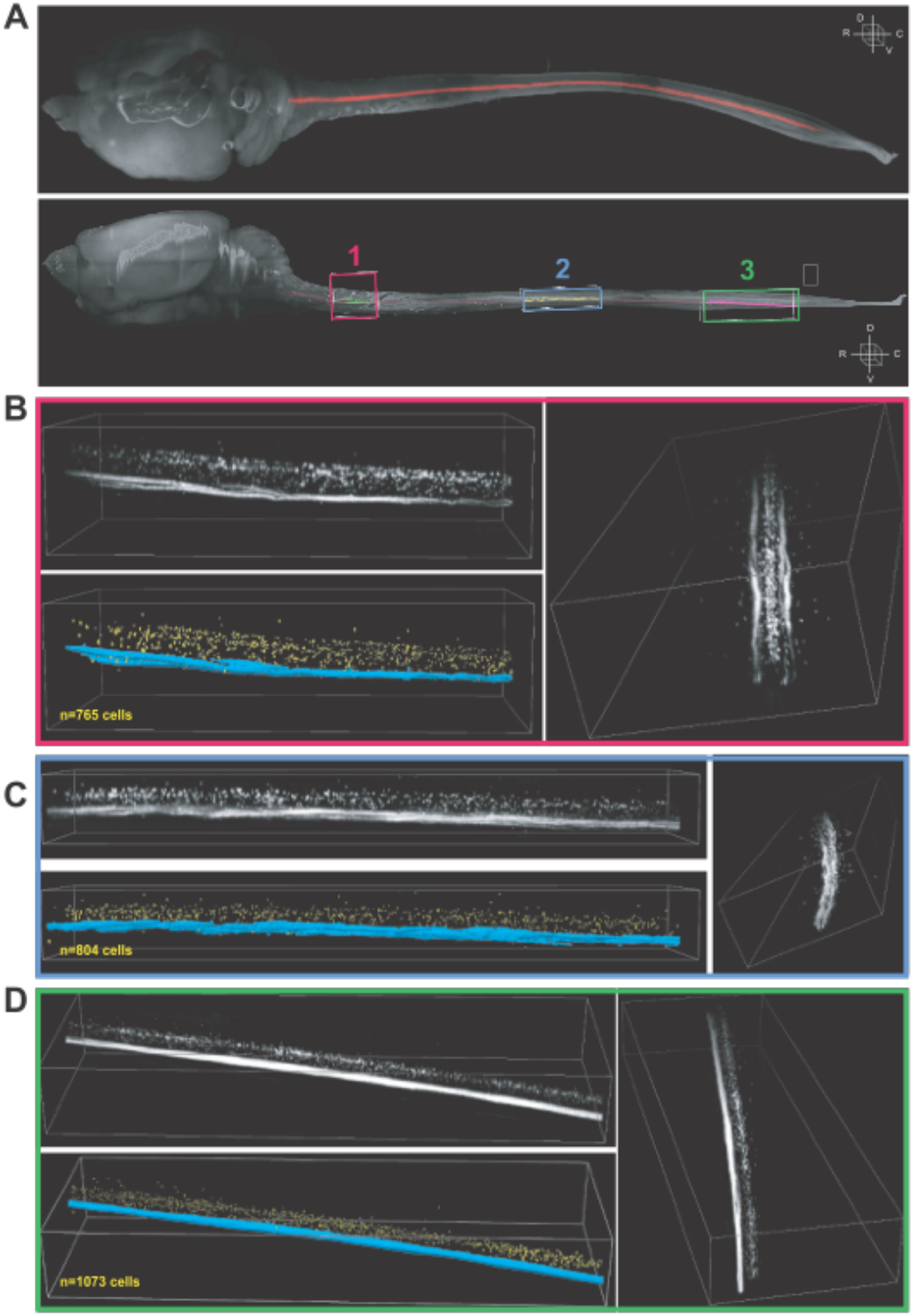
Localization of CSF-contacting neurons in the mouse Central Nervous System. **A)** Top. Dorsal view of the full rostro-caudal distribution of the CSF-cN system (red) within the mouse CNS (grey). Bottom. Lateral view of the image shown in the Top panel. Three boxes are indicated to depict representative cervical (1, red), thoracic (2, blue), and lumbar (3, green) regions acquired at higher magnifications. Boxes sizes are in μm along the Rostro-caudal (RC) x Latero-Median (LM) x Dorso-Ventral (DV): (1) 1764.75 × 1825.50 × 1726.00; (2) 3076.13 × 1784.25 × 818.00; (3) 3734.25 × 1228.50 × 1192.00. **B-D)** Higher magnification views and segmentation for the cervical (B), thoracic (C) and lumbar (D) SC segments for the boxes labelled in panel 1A Bottom as 1, 2 or 3 respectively (see color code in A). For each panel: Top Left, Higher magnification views of the respective boxes illustrated in **Fig. 1A, Bottom** panel with the same orientation. **Bottom Left**. 3D representation of the segmentation and cell counting of CSF-cNs for each SC segments (see methods; CSF-cNs in yellow and axon bundles in blue). **Right**. 90° rotation view of the image in the Top Left panel. Dimensions of the box are for the RC × LM × DV axes (in μm): cervical segment (B), 1764.75 × 913.25 × 463.00; thoracic segment (B), 3076.13 × 658.15 × 322.00 and lumbar segment (D), 3734.25 × 1228.50 × 492.00.

**Figure 2.**
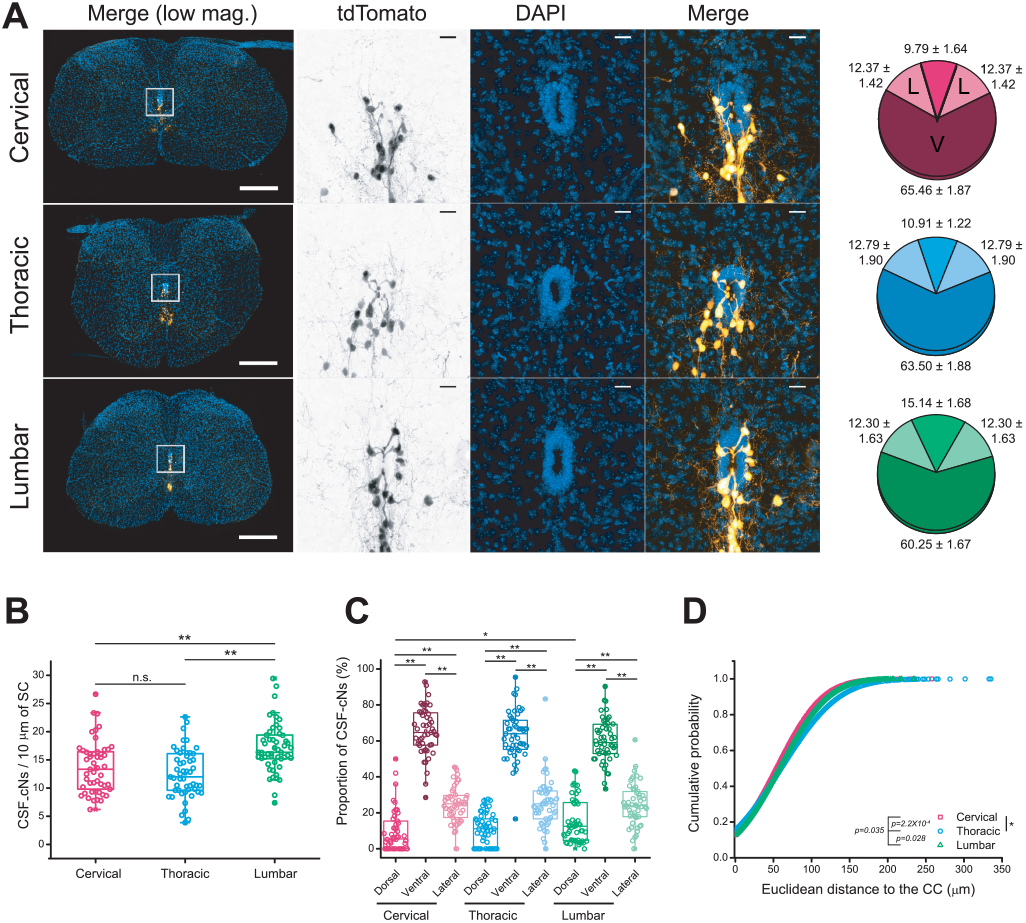
Confocal analysis of CSF-cNs distribution in the mouse spinal cord. **A) Left**. Confocal images of tdTomato fluorescence in 3 weeks old PKD-tdTomato mouse at cervical (**Top** panels), thoracic (**Middle** panels), and lumbar (**Bottom** panels) levels of the SC. tdTomato fluorescence is depicted as inverted greyscale image when presented alone, and as an Orange hot lookup table in merge images. DAPI is depicted as Azure lookup table. Scale bar=200 μm in **Left** panels; 20 μm in Middle and Right panels. **Right**. Pie plots of CSF-cN distribution in dorsal, lateral, and ventral quadrants at cervical (**Top**, red), thoracic (**Middle**, blue) and lumbar (**Bottom**, green) levels of the SC. **B)** Density of CSF-cNs per 10 μm of SC at cervical (red), thoracic (blue), and lumbar (green) levels of the SC. **C)** CSF-cNs distribution at the different levels of the SC. The color code corresponds to the one depicted in figure 1A. Kruskal-Wallis ANOVA followed by Dunn’s test. *p<0.05; **p<0.01. For the sake of clarity, only significant comparisons are indicated. **D)** Cumulative distribution function plot of the Euclidean distances of CSF-cNs to the lumen of the CC at cervical (red), thoracic (blue), and lumbar (green) levels of the SC. Kolmogorov-Smirnov test. ***p<0.001.

These findings suggest that, along the SC, CSF-cNs form a homogenous population with a conserved morphology and a predominantly ventral localization and a larger density in the lumbar region.

### Along the rostro-caudal axis CSF-cNs exhibit similar intrinsic properties

#### CSF-cNs intrinsic properties and firing pattern

CSF-cNs are found along the entire CC axis. However, only medullar CSF-cNs in mice have been extensively studied^12,22^. Here we characterized the intrinsic properties of lumbar (L-), thoracic (T-), and cervical (C-) CSF-cNs to determine if they form a homogenous population. Consistent with previous studies in the brainstem^12,22^, spinal CSF-cNs have a high input resistance (R_*m*_, 3.6±2.3 GΩ; N=191, n=659) and small membrane capacitance (C_*m*_, 5.5±2.8 pF; N=191, n=659), with a fast membrane time constant (τ_*m*_, 0.09±0.05 ms; N=191, n=659) in agreement with their small soma and dendritic arborization. When comparing across spinal segments, L-CSF-cNs displayed a higher R_*m*_, while T-CSF-cNs had the lowest C_*m*_ (***Table 1***).

**Table 1.**
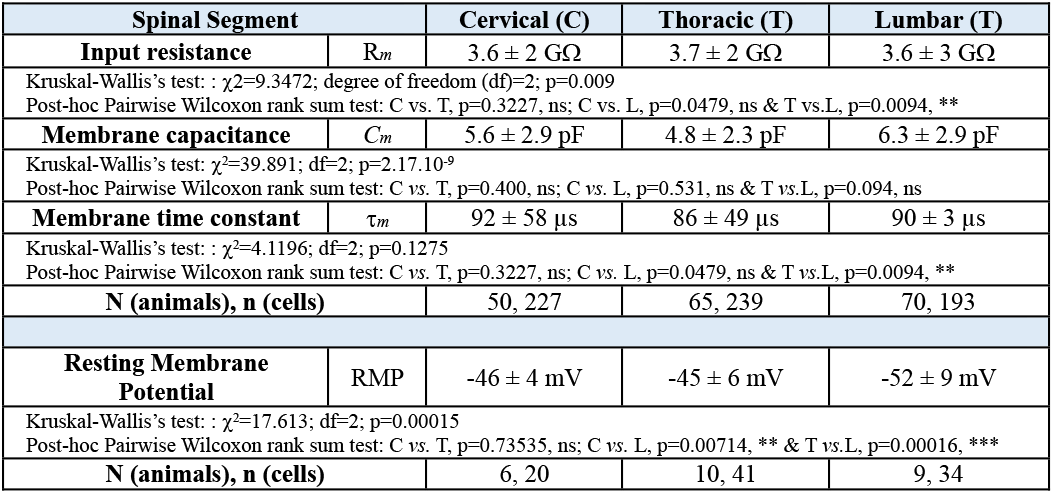
Intrinsic properties of CSF-cNs along the rostro-caudal central canal axis.

In current-clamp mode (CC, I=0), resting membrane potentials (RMP) is on average -55±19 mV (N=25, n=95), with L-CSF-cNs being more hyperpolarized compared to more depolarized CSF-cNs in anterior regions (***Table 1***; aCSF and intracellular solution A, ***Supplementary Tables 1 and 2***). AP discharge patterns were also assessed. In line with previous findings in juvenile rats^9^ and postnatal mice^8^, spinal CSF-cNs exhibit either tonic or single spike discharges. Following a current injection step (+20 pA, 200-500 ms duration and membrane potential at -60 mV, DC current: -10 to -15 pA), 58% of the neurons showed tonic AP firing (38 cells out of 65 recorded), while 42% fired a single AP (27 cells out of 65 recorded; ***Table 2***). Comparative analysis revealed that most L-CSF-cNs and C-CSF-cNs had single AP discharges while T-CSF-cNs exhibited a primarily tonic pattern (***Table 2***). However, these differences were not statistically significant and did not depend on the neuron’s position around the CC. Overall, the findings suggest that while CSF-cNs share similar properties and appear to form a largely homogenous population throughout the SC, although some regional exist.

**Table 2.**
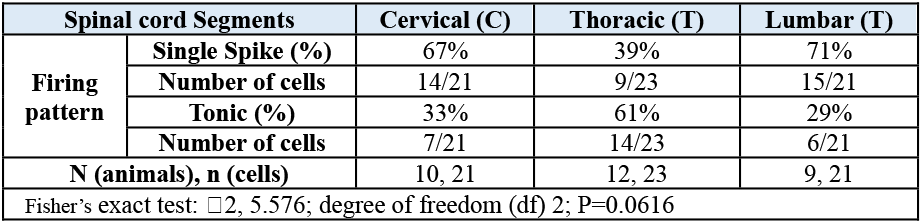
CSF-cNs exhibit either a single spike or tonic AP discharge pattern.

#### CSF-cN modulation by extracellular pH through the activity of PKD2L1 and ASICs

In agreement with their expression of PKD2L1 channels^5,12,22,23,28^, we observed spontaneous unitary PKD2L1 activity in CSF-cNs recorded at V_*h*_ -80 mV (aCSF with all synaptic receptors blocked, see *Material and Methods*), with an average current amplitude of -13±2 pA and an open probability (NPo) of 0.03±0.04 (N=23, n=60; ***Supplementary Figure 3***). No differences were found in amplitude or NPo across spinal segments. PKD2L1, a sensory chemoreceptor, responds to pH changes^12,22,23^. Exposure to alkaline pH (pH 9; ***Supplementary Figure 3A-D***) causes no change in current amplitude (-13±2, - 13±2 and -13±2 pA in control, pH 9 and Wash, respectively; N=13, n=32; *Kruskal-Wallis rank sum test*: 𝒳^2^=2.7856, df=2, p(𝒳^2^)=0.2484 and a posthoc *Pairwise comparisons using Wilcoxon rank sum test* with continuity correction: CTR *vs*. pH 9, p=0.36; CTR *vs*. Wash, p=0.36 and Wash *vs*. pH 9, p=0.57) but increased NPo by 20% (16±30%) from 0.04±0.05 to 0.16±0.12 before returning to baseline value upon Wash (0.08±0.1; *Kruskal-Wallis rank sum test*: 𝒳^2^=27.219, df=2, p(𝒳^2^)=1.229.10^−6^ and a posthoc *Pairwise comparisons using Wilcoxon rank sum test* with continuity correction: CTR *vs*. pH 9, p=1.3.10^−7^; CTR *vs*. Wash, p=0.16162 and Wash *vs*. pH 9, p=0.00045; N=13, n=32). Again, all spinal CSF-cNs show a similar response to alkaline pH exposure (***Supplementary Figure 3A-B***).

PKD2L1 acts as a AP generator at the single channel level and exposure to alkaline pH increases PKD2L1 activity as well as CSF-cN excitability in the *medulla*^22^. In recordings conducted in current-clamp mode at RMP (aCSF with all synaptic receptors blocked, see *Material and Methods*), all spinal CSF-cNs exhibit an increase in the AP firing frequency from 1.6±2.1 Hz in control to 2.4±2.4 Hz under alkaline pH, which also reversed upon Wash (frequency in Wash: 1.4±2.0 Hz; ***Supplementary Figure 3C, D***; *Kruskal-Wallis rank sum test*: 𝒳^2^=9.4037, df=2, p(𝒳^2^)= 0.009079 and a posthoc *Pairwise comparisons using Wilcoxon rank sum test* with continuity correction: CTR *vs*. pH 9, p=0.012; CTR *vs*. Wash, p=0.902 and Wash *vs*. pH 9, p=0.012; N=12, n=32).

In contrast, exposure to acidic pH (pH 5; same extracellular and intracellular solutions as above) inhibited PKD2L1 activity, reducing the current amplitude from -13±2 pA in control to 0.9±2.1 pA in pH5 (***Supplementary Figure 3E***), followed by a recovery to -14±2 pA upon Wash (N=10, n=28; *Kruskal-Wallis rank sum test*: 𝒳^2^=55.896, df=2, p(𝒳^2^)= 7.282.10^−13^ and a posthoc *Pairwise comparisons using Wilcoxon rank sum test* with continuity correction: CTR *vs*. pH5, p=3.9.10^−16^; CTR *vs*. Wash, p=0.27 and Wash *vs*. pH5, p=3.9.10^−16^). NPo also decreased reversibly under acidic conditions from 0.024±0.018 in control to 0.001±0.001 in the presence of acidic solution (***Supplementary Figure 3E,F***; Wash: 0.048±0.076; N=10 n=28; *Kruskal-Wallis rank sum test*: 𝒳^2^=52.84, df=2, p(𝒳^2^)=3.357.10^−12^ and a posthoc *Pairwise comparisons using Wilcoxon rank sum test* with continuity correction: CTR *vs*. pH5, p=7.1.10^−10^; CTR *vs*. Wash, p=0.95 and Wash *vs*. pH5, p=8.2.10^−10^).

Additionally, acidic pH triggered in all spinal CSF-cNs a large transient inward current (on average -801±411 pA; N=10, n=24) followed by a persistent phase (***Supplementary Figure 3G***), typical of ASICs activation and previously reported in medullar CSF-cNs^12,22^ (see also Jalavand and Colleagues^24^). The ASICs currents had similar amplitude in all spinal CSF-cNs recorded (C: - 712±289 pA; T: -685±189 pA and L: -1005±599 pA (n=8 for each segments); *Kruskal-Wallis rank sum test*: 𝒳^2^=0.74, df=2, p(𝒳^2^)=0.6907 and a posthoc *Pairwise comparisons using Wilcoxon rank sum test* with continuity correction: C *vs*. T, p=0.76; C *vs*. L, p=0.96 and T *vs*. L, p=0.76). During the persistent phase of ASICs current, PKD2L1 channels were inhibited (colored boxes in ***Supplementary Figure 3G*** and above). In current-clamp recordings at RMP, ASICs activation led on average to a +37±9 mV depolarization (***Supplementary Figure 3H, I***_***1***_; N=11, n=38) that was large enough to trigger APs in all spinal CSF-cNs (***Supplementary Figure 3H, I***_***2***_).

In conclusion, spinal CSF-cNs maintain conserved intrinsic properties with a depolarized membrane potential and exhibit chemosensory functions, responding to extracellular pH variations.

### Spinal CSF-cNs express functional sodium, potassium and calcium voltage-dependent channels

Marichal and colleagues^9^ showed functional sodium (Na_V_) and potassium (K_V_) voltage-dependent conductances in juvenile rat CSF-cNs. We analyzed mouse spinal CSF-cNs to identify the presence and types of voltage-dependent conductances.

We first analyzed Na^+^ voltage-dependent currents. Recordings of spinal CSF-cNs in voltage-clamp mode at V_*h*_ -70 mV with specific solutions to isolate I_Na_ currents (Na_V_ solution and intracellular solution B, ***Supplementary Tables 1 and 2***) reveals fast, inactivating inward currents in all cells during incremental voltage steps (V_*Step*_ from -70 to +60 mV, ΔV=+10 mV, 100 ms; ***Fig. 3A***). The current activates at V_*Step*_ more depolarized than -40 mV, peaks at 0 mV with an average current amplitude of -712±450 pA (N=10, n=41) and a current density of -124±67 pA.pF^−1^ (N=10, n=41; ***Fig. 3B, C***). Subsequently the current decreases for more depolarized V_*Step*_ and reversed above +50 mV in agreement with the calculated sodium equilibrium potential (E_Na_=+46 mV, ***Supplementary Table 1***). We did not observe difference in the Na^+^ current amplitude between C-, T- and L-CSF-cNs (***Fig. 3C***). Tetrodotoxine (TTX, 0.5 µM) completely blocked the current confirming expression of functional TTX-sensitive Na_V_ channels in spinal CSF-cNs (***Fig. 3D***; inhibition by 106±9% from -141±71 in control to 8±11 pA.pF^-1^ in TTX (N=10, n=28); cervical, red: -139±65 pA.pF^-^ and 8±11, pA.pF^−1^ (N=4, n=12); thoracic, blue: −178±92 pA.pF^−1^ and 10±14 pA.pF^−1^ (N=4, n=6) and lumbar, green: −122±61 pA.pF^−1^ and 7±9 pA.pF^−1^ (N=2, n=10) in control and TTX, respectively; *ANOVA*.*lme*: F=25.87, df=5 and 50, p(F)=8.582.10^−13^ and *Tukey (EMM) post-hoc* test to compare CTR *vs*. TTX within the different regions: ***,p<0.0001 in cervical, thoracic and lumbar segments. There is no difference between CTR and TTX between regions).

**Figure 3.**
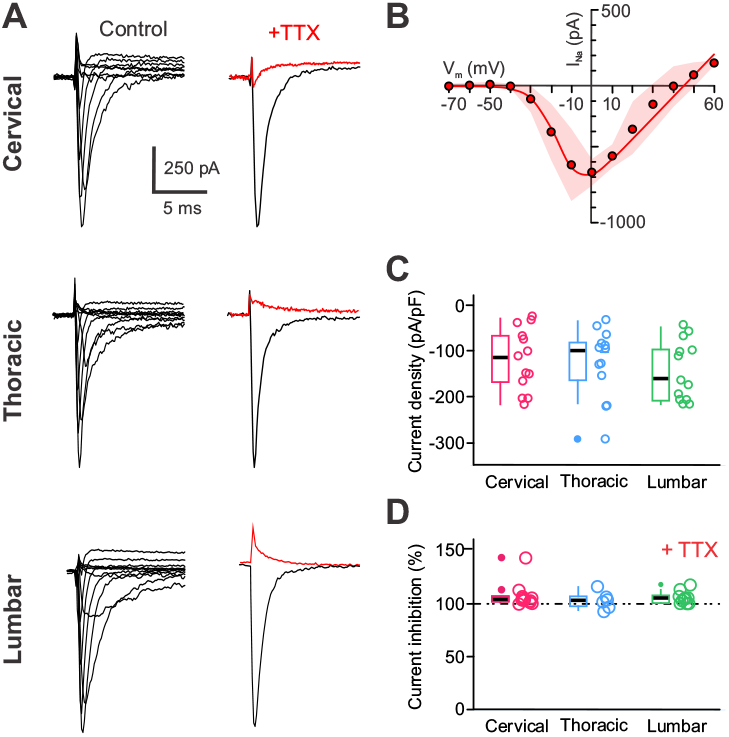
CSF-cNs express functional TTX-sensitive sodium voltage-dependent channels. **A)** Left. Representative whole-cell current traces recorded in response to V_Step_ from -70 mV to +60 mV (+10 mV increments for 50 ms) from V_h_ -80 mV to elicit Na^+^ current in a CSF-cN at the cervical (Top), thoracic (Middle) and lumbar (Bottom) levels. Right. Na^+^ whole-cell currents elicited with a voltage step to +10 mV from V_h_ -70 mV in control (black trace) and in the presence of 0.5 µM TTX (red traces) recorded in the CSF-cNs illustrated Left. **B)** Average current-voltage relationship (IV-curve) for the Na^+^ currents recorded in CSF-cNs at all levels and generated from the peak current amplitude measured for each V_Step_ and presented against the respective V_Step_. **C)** Summary box-and-whiskers plots of the averaged amplitude of the Na^+^ peak current density recorded in control at a V_Step_ of +10 mV from V_h_ -80 mV from CSF-cNs at each segment of interest (cervical, red: -132±74 pA.pF^-1^ (N=8, n=23); thoracic, blue: -91±40 pA.pF^-1^ (N=8, n=13) and lumbar, green: -129±63 pA.pF^-1^ (N=4, n=20); (Kruskal-Wallis rank sum test: χ2=2.2544, df=2, p=0.3239; posthoc Pairwise comparisons using Wilcoxon rank sum test with continuity correction: C vs. L, p=0.88, T vs. C and T vs. L p=0.29). **D)** Summary box-and-whiskers plots of the averaged TTX inhibition (in percent) of the Na^+^ current inhibition by 108±12 %, 103±8 % and 106±6 % for cervical (N=4, n=12), thoracic (N=4, n=8) and lumbar (N=2, n=6), respectively; Kruskal-Wallis rank sum test: 𝒳^2^=0.65025, df=2, p(𝒳^2^)=0.7224 and a post hoc Pairwise comparisons using Wilcoxon rank sum test with continuity correction: C vs. T, p=0.83; C vs. L, p=0.87 and T vs. L, p=0.83). In **C** and **D**, Single data points for all the recorded cells at each level are presented with colored opened circle (same color code as above).

In juvenile rats9, delayed rectifier (I_KD_) and A-type K^+^ currents (I_A_) were identified. To determine the functional expression of voltage-dependent K^+^ channels (K_V_) in mouse spinal CSF-cNs, we conducted recordings at V_*h*_ −80 mV using specific solutions to isolate K_V_ currents (K_V_ solution and intracellular solution A, ***Supplementary Tables 1 and 2***) and applied a 50 ms pre-step to - 100 mV, followed by V_*Step*_ from −40 to +60 mV (ΔV=+10 mV, 100 ms; ***Fig. 4A***). The elicited current had an average current amplitude at the peak of 5478±2012 pA and a current density of 900±402 pA.pF^−1^ at +60 mV (N=14, n=60). The I–V curve (***Fig. 4B***), in agreement with the potassium reversal potential (E_K_=−94 mV, ***Supplementary Table 1***), shows outward currents activated for V_*Step*_ larger than −30 mV and the current increased proportionally to the V_*Step*_ amplitude. Typically, the activated currents presented an initial transient phase (Peak (E, *): 900±402 pA.pF^−1^ followed by a persistent one (Persistent (P, #): 778±368 pA.pF^−1^; N=14, n=60) lasting for the whole duration of V_*Step*_.

**Figure 4.**
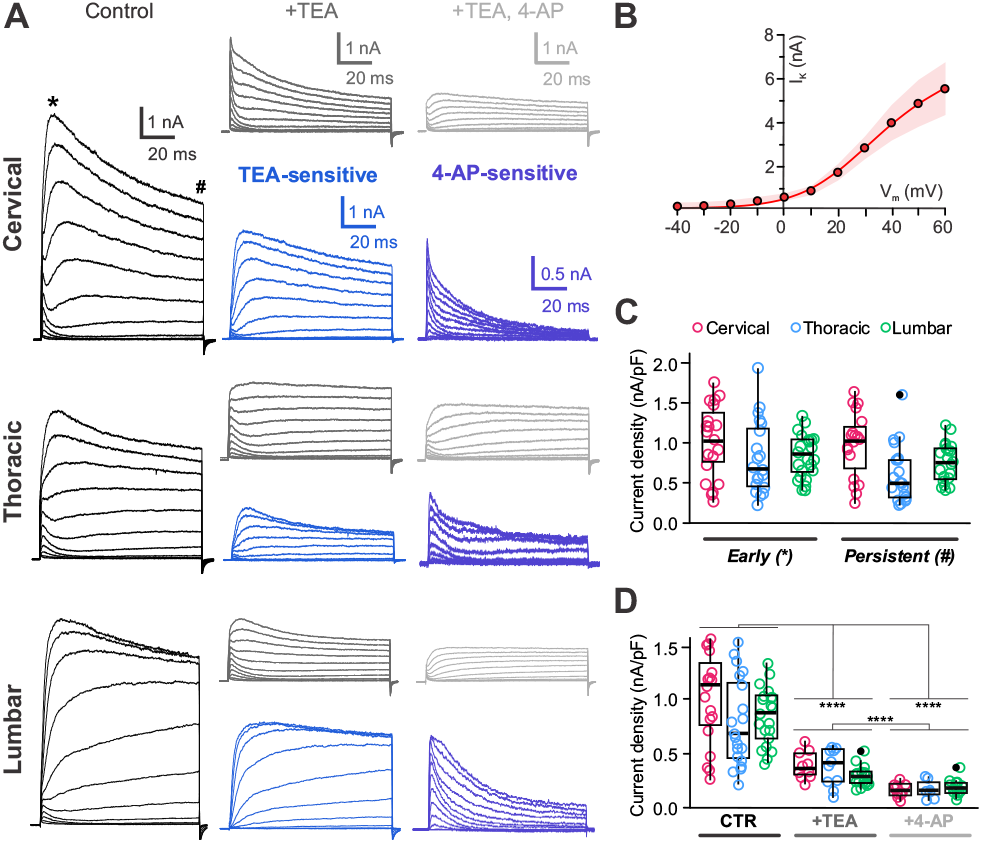
Delayed rectifier and A-type voltage-dependent potassium channels are present in CSF-cNs. **A)** Representative whole-cell current traces recorded in response to V_Step_ from -40 to +60 mV (increments of 10 mV, 100 ms) from V_h_ -80 mV in control (black traces), in the presence of TEA (grey traces) and in the presence of TEA and 4-AP (light grey traces). cervical, thoracic and lumbar levels, from Top to Bottom. For each level the TEA-(Blue traces) and 4-AP-senstive (violet) currents are presented (see Results for more details). **B)** Control average current-voltage relationship (IV-curve) for the K^+^ currents recorded in CSF-cNs from all levels. **C)** Summary box-and-whiskers plots of the averaged amplitude of the peak K^+^ current density recorded in control (V_Step_ +60 mV from V_h_ -80 mV) for the peak/early (E,*) and persistent (P, #) currents (cervical, red: 1039±450 and 976±408 pA.pF^-1^ (N=4, n=19); thoracic, blue: 814±453 and 612±364 pA.pF^-1^ (N=4, n=21) and lumbar, green: 855±267 and 758±243 pA.pF^-1^ (N=2, n=20) - E and P in order; ANOVA.lme: F=3.35, df=5 and 114, p(F)=0.007349 and Tukey (EMM) post-hoc test to compare E vs. P within the different regions and between regions; * and ** for p<0.05 and <0.01, respectively). **D)** Summary box-and-whiskers plots of the averaged amplitude of the peak K^+^ current density for CSF-cNs recorded (V_Step_ +60 mV from V_h_ -80 mV) in the 3 regions of interest in control (CTR), in the presence of TEA (+TEA) and of TEA + 4-AP (+4-AP). (C, red: 1039±450, 397±128168±64 pA.pF^-1^ (N=4, n=19, 10, 10); T, blue: 814±453, 380±178 and 176±87 pA.pF^-1^ (N=5, n=20,10, 7) and L, green: 855±267, 296±88 and 196±67 pA.pF^-1^ (N=5, n=21, 19, 17); data given for CTR, +TEA and +4-AP, respectively; ANOVA.lme: F=22.73, df=8 and 124, p(F)<2.2.10^−16^ and Tukey (EMM) post-hoc test to compare CTR vs. +TEA, CTR vs. +4-AP and +TEA vs. +4-AP within the different regions and between the different regions; *, ** and *** for p<0.05, <0.01 and <0.001, respectively). In C and D, Single data points for all the recorded cells at each level are presented with colored opened circle (same color code as above).

Application of TEA (10 mM) reduced the current by 56±18% (current density of 344±131 pA.pF^−1^; N=14, n=39; ***Fig. 4A***, current traces +TEA and ***Fig. 4D***) and the TEA-sensitive (digital subtraction of the current recorded in control minus that in TEA; blue traces in ***Fig. 4A***) exhibited characteristic kinetics of I_KD_ (***Fig 4A***). Adding 4-AP (4 mM; +4-AP) further decreased the current by 44±29% (amplitude of 183±69 pA.pF^−1^; N=14, n=34; ***Fig 4A and D***). The 4-AP sensitive current (digital subtraction of the current recorded in TEA only minus that in TEA and 4-AP; violet traces in ***Fig*.*4A***) exhibit fast-rising phase, and rapid inactivation compatible with I_A_ (***Fig. 4A***). The remaining current was likely non-selective or K^+^ leak currents.

In a previous report, we indicated that mouse medullar CSF-cNs express functional voltage-dependent Ca^2+^ channel (Ca_V_) mainly of the N-type (Ca_V_2.2)^19^ while in juvenile rat^9^ they appear to express both High (HVA) and Low Voltage-Activated (LVA) Ca^2+^ conductances. To investigate this in spinal CSF-cNs, we isolated Ca^2+^ currents (Ca_V_ solution and intracellular solution B, ***Supplementary Tables 1 and 2***) and recorded neurons at V_*h*_ −80 mV with V_*Step*_ from −40 to +30 mV (ΔV=+10 mV, 100 ms). The elicited inward currents show a fast rise, an initial transient decrease followed by a persistent steady-state current. They also exhibit fast tail currents upon repolarization of the membrane potential (***Fig. 5A***). At V_*h*_ −60 mV, the I-V curve revealed that currents activate at V_*Step*_ larger than −40 mV, peaks at 0 mV (***Fig. 5A, B***), with an average current amplitude of −278±168 pA and current density of −48±34 pA.pF^−1^ (N=17, n=58; ***Fig. 5C***). T-CSF-cNs showed the largest Ca^2+^ current peak amplitude while it was the smallest in C-CSF-cNs (***Fig. 5C***). Altogether the properties of the currents recorded agree with those mediated by Ca_V_ and this observation was further confirmed by a current block of 112±10 % from −46±39 pA.pF^−1^ in control to 4±4 pA.pF^−1^ in the presence of Cd^2+^ (200 µM, pressure application; ***Fig. 5A, Right and D***; N=17, n=29), a Ca_V_ selective blocker (. In juvenile rat^9^ and mouse thoracic segment, CSF-cNs were suggested to also express LVA Ca_V_ of the T-type (Ca_V_ 3.3) that are inactivated for membrane potential around −60 mV. Using ramp protocols from V_*h*_ −80 mV, revealed a characteristic ‘shoulder’ in all spinal CSF-cNs (average amplitude: −51±32 pA, membrane potential: −29±7 mV (N=19, n=172); ***Fig. 5E***). This shoulder was not observed in recordings carried out at V_*h*_ −60 mV, a membrane potential where T-type Ca^2+^ (Ca_V_ 3) are known to be inactivated (***Fig. 5E***). Note that the current traces illustrated in ***Figure 5E*** have been normalized to the peak value for a better visualization. These results demonstrate that spinal CSF-cNs express both HVA and LVA Ca_V_, with T-CSF-cNs having higher Ca^2+^ current density.

**Figure 5.**
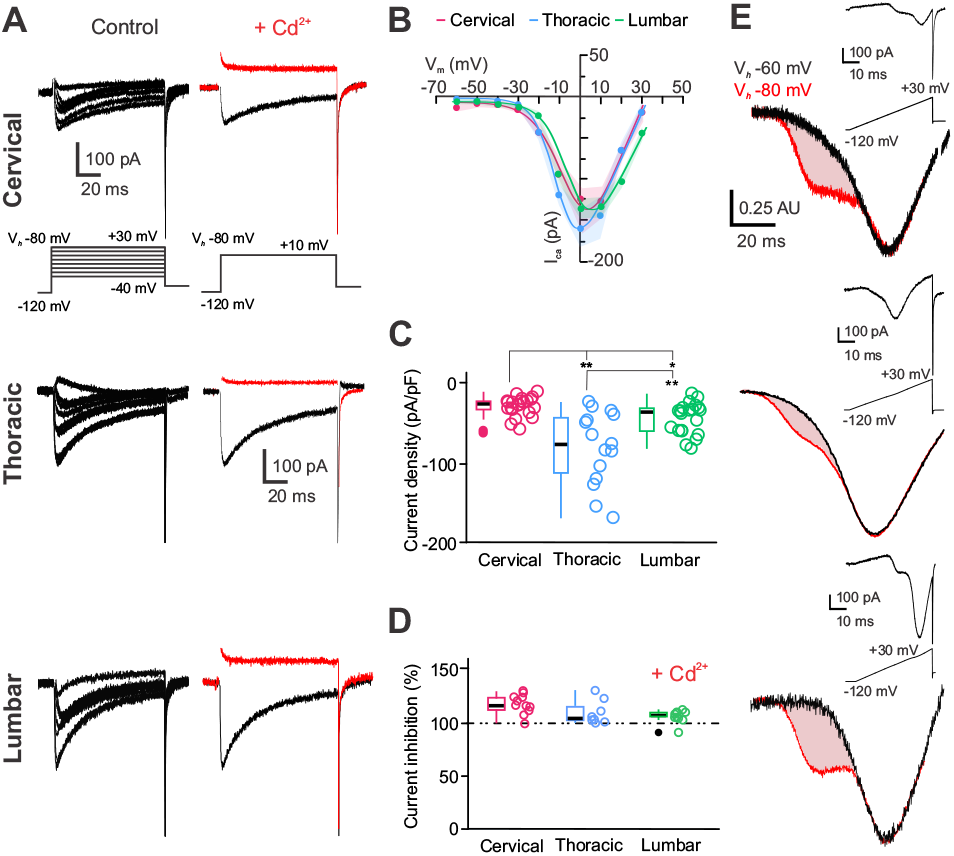
CSF-cNs express calcium channels of the high and low voltage-activated types. **A)** Left. Representative whole-cell current traces recorded in response to V_Step_ from V_Step_ -40 mV to +30 mV (V_Step_, 10 mV increments, 100 ms) from V_h_ -80 mV with a prepulse to -120 mV to elicit I_Ca_ in a CSF-cN at the cervical (Top), thoracic (Middle) and lumbar (Bottom) levels. Right. Ca^2+^ whole-cell currents elicited with a voltage step to +10 mV from V_h_ - 80 mV in control (black trace) and in the presence of 200 µM Cd^2+^ (red traces) recorded in the CSF-cNs illustrated Left. **B)** Average current-voltage relationship (IV-curve) for the I_Ca_ recorded from V_h_ -60 mV in CSF-cNs at the cervical (red, N=10, n=41), thoracic (blue, N=10, n=41) and lumbar (green, N=10, n=41) using a ramp protocol for current against time and subsequently converted as current against voltage relationship. **C)** Summary box-and-whiskers plots of the averaged amplitude of the Ca^2+^ peak current density recorded in control at a V_Step_ of +10 mV from V_h_ -80 mV and measured at the steady state in CSF-cNs from each segment of interest (cervical, red: -29±12 pA.pF^-1^ (N=2, n=22); thoracic, blue: -81±46 pA.pF^-1^ (N=5, n=15) and lumbar, green: -44±20 pA.pF^-1^ (N=10, n=21); Kruskal-Wallis rank sum test: 𝒳^2^=20.143, df=2, p(𝒳^2^)=4.226.10^−5^ and a post hoc Pairwise comparisons using Wilcoxon rank sum test with continuity correction: C vs. T, p=5.6.10^−5^; C vs. L, p=0.0049 and T vs. L, p=0.0110). **D)** Summary box-and-whiskers plots of the averaged Cd^2+^ inhibition (in percent) of I_Ca_ (inhibition by 118±9 % for C-(N=2, n=11); 110±11 % for T-(N=5, n=8) and 106±6 % for L-CSF-cNs (N=10 n=10), respectively; Kruskal-Wallis rank sum test: 𝒳^2^=0.38571, df=2, p(𝒳^2^)=0.8246 and a post hoc Pairwise comparisons using Wilcoxon rank sum test with continuity correction: C vs. T, C vs. L and T vs. L, p=0.97, respectively). In **C** and **D**, Single data points for all the recorded cells at each level are presented with colored opened circle (same color code as above). **E)** Averaged Ca^2+^ IV-Curves converted the recordings obtained using a ramp protocol from V_h_ -80 mV (red traces) and -60 mV (black traces) for CSF-cNs at the cervical to the lumbar levels (Top to Bottom; see Methods for more details). The red shaded area indicates the LVA (T-type channel) currents. Note that the data are normalized to the peak and presented in arbitrary units (AU) for a better comparison. Inset illustrates the activation protocol and an elicited representative recording for each region of interest.

Altogether, spinal CSF-cNs express functional Na_V_, along with K_V_ of the delayed rectifier-(I_KD_) and A-type (I_A_), which are similarly distributed along the CC axis and would be responsible for the AP generation. Additionally, all spinal CSF-cNs express both HVA and LVA Ca_V_, with higher current density in T-CSF-cNs.

### CSF-cN express the major ionotropic synaptic receptors

Mice medullar CSF-cNs express synaptic ionotropic receptors, including GABAergic, glycinergic, and AMPA/Kainate glutamatergic receptors^12^,mediating spontaneous and electrically evoked synaptic responses^19^. To assess the expression of these receptors in spinal CSF-cNs, we performed voltage-clamp recordings at V_*h*_ −80 mV and applied selective agonists by pressure.

GABA (1 mM, 30 ms; ***Fig. 6A***) and glycine (1 mM, 100 ms; ***Fig. 6C***) elicited fast and large inward currents (with E_Cl_=+5 mV, aCSF and intracellular solution C, ***Supplementary Tables 1 and 2***) in all CSF-cNs. The average current densities of -321±36 pA.pF^−1^ (N=6, n=36) and -529±64 pA.pF^−1^ (N=8, n=32) for GABA and glycine application, respectively (***Fig. 6B, D***). The currents were blocked in the presence of their selective antagonist, gabazine (Gbz,10 µM; 97.6±0.3 %; N=6, n=27) and strychnine (Stry,1 µM; 95.5±1.0 %; N=9, n=23), respectively and therefore mediated by GABA_A_ and glycine receptor activation (***Fig. 6B***, red traces). The current recorded were mediated by GABA and glycine receptor-mediated currents were blocked by gabazine (97.6±0.3%; N=6, n=27) and strychnine (95.5±1.0%; N=9, n=23). GABA currents were highest in T-CSF-cNs, while glycine currents were larger in L-CSF-cNs. When comparing the data along the CC axis, GABA_A_-mediated currents were the highest in T-CSF-cNs compared to C- and L-CSF-cNs (***Fig. 6A, B***). In contrast, glycine-mediated current where similar in C- and T-CSF-but larger in L-CSF-cNs (***Fig. 6C, D***).

**Figure 6.**
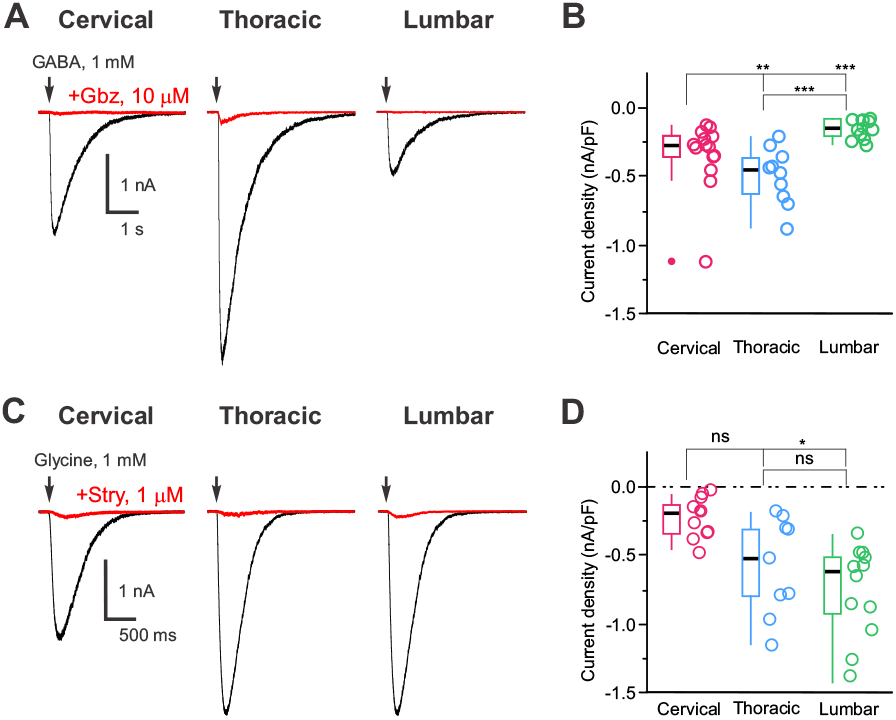
CSF-cNs express different density of GABA_A_ and glycine receptors. **A)** Representative whole-cell current traces recorded at V_h_ -80 mV in response to pressure application of GABA (1 mM, 30 ms) in control (black traces) and in the presence of gabazine (Gbz,10 µM; red traces), a selective GABA_A_ receptor antagonist, in CSF-cNs from the cervical, thoracic and lumbar segments (**Left to Right**). **B)** Summary box-and-whiskers plots of the averaged amplitude of the GABA-mediated current densities recorded in control at V_h_ -80 mV (cervical, red: -342±250 pA.pF^-1^, N=2, n=14; thoracic, blue: -495±204 pA.pF^-1^, N=2, n=10 and lumbar, green: -153±69 pA.pF^-1^, N=2, n=12; Kruskal-Wallis rank sum test: 𝒳^2^=17.975, df=2, p(𝒳^2^)=1.25.10^−4^ and a post hoc Pairwise comparisons using Wilcoxon rank sum test with continuity correction: C vs. T, p=0.02602; C vs. L, p=0.00306 and T vs. L, p=0.00011). **C)** Representative whole-cell current traces recorded at V_h_ -80 mV in response to pressure application of glycine (1 mM, 30 ms) in control (black traces) and in the presence of strychnine (Stry,10 µM; red traces), a selective glycine receptor antagonist, in CSF-cNs from the cervical, thoracic and lumbar segments (**Left to Right**). **D)** Summary box-and-whiskers plots of the averaged amplitude of the glycine-mediated current densities recorded in control at V_h_ -80 mV (cervical, red: -232±146 pA.pF^-1^, N=2, n=11; thoracic, blue: -584±351 pA.pF^-1^, N=9, n=10 and lumbar, green: -761±340 pA.pF^-1^, N=6, n=12; Kruskal-Wallis rank sum test: 𝒳^2^=14.979, df=2, p(𝒳^2^)=5.589.10^−4^ and a post hoc Pairwise comparisons using Wilcoxon rank sum test with continuity correction: C vs. T, p=0.03; C vs. L, p=5.3e-05 and T vs. L, p=0.25). In **B** and **D**, Single data points for all the recorded cells at each level are presented with colored opened circle (same color code as above).

Next, medullar CSF-cNs express functional AMPA/Kainate receptors^12,22^, and our results show that pressure application of glutamate (100 µM, 100-500 ms) induces inward currents in all spinal CSF-cNs with an average current amplitude and density of -22±14 pA and -6±5 pA.pF^-1^ and characterized by slow kinetics similar to those observed in medullar CSF-cNs (***Fig. 7A***_***1***_ (N=22, n=46), *Kruskal-Wallis rank sum* test: 𝒳^2^=32.74, df=2, p(𝒳^2^)= 7.773.10^−8^ and a post hoc *Pairwise comparisons using Wilcoxon rank sum test with continuity correction*: C *vs*. T, p=0.0015; C *vs*. L and T *vs*. L, p<0.0001). These currents were present across spinal segments but were significantly smaller in L-CSF-cNs (***Fig. 7B***) (aCSF and intracellular solution C, ***Supplementary Tables 1 and 2***). AMPA/Kainate receptor activation was confirmed by the selective current inhibition in the presence of DNQX (400 µM; 91±2% current inhibition). AMPA (100 µM, ***Fig. 7A***_***2***_) and Kainate (100 µM, ***Fig. 7A***_***3***_) produced similar responses, while NMDA (100 µM) did not induce currents, suggesting the expression of AMPA/Kainate but not NMDA receptors in spinal CSF-cNs. In current-clamp mode at -60 mV (DC current injection of -10 to -15 pA; (aCSF and intracellular solution A, ***Supplementary Tables 1 and 2***), glutamate application caused significant depolarization (+36±6.5 mV, ***Fig. 7D, E***_***1***_; N=4, n=21), with larger depolarizations in C- and T-CSF-cNs than L-CSF-cNs (***Fig. 7E***_***1***_). This depolarization led to AP firing in C- and T-CSF-cNs but not in L-CSF-cNs (***Fig. 7E***_***2***_), and was absent with DNQX blockade (***Fig. 7D***, red traces).

**Figure 7.**
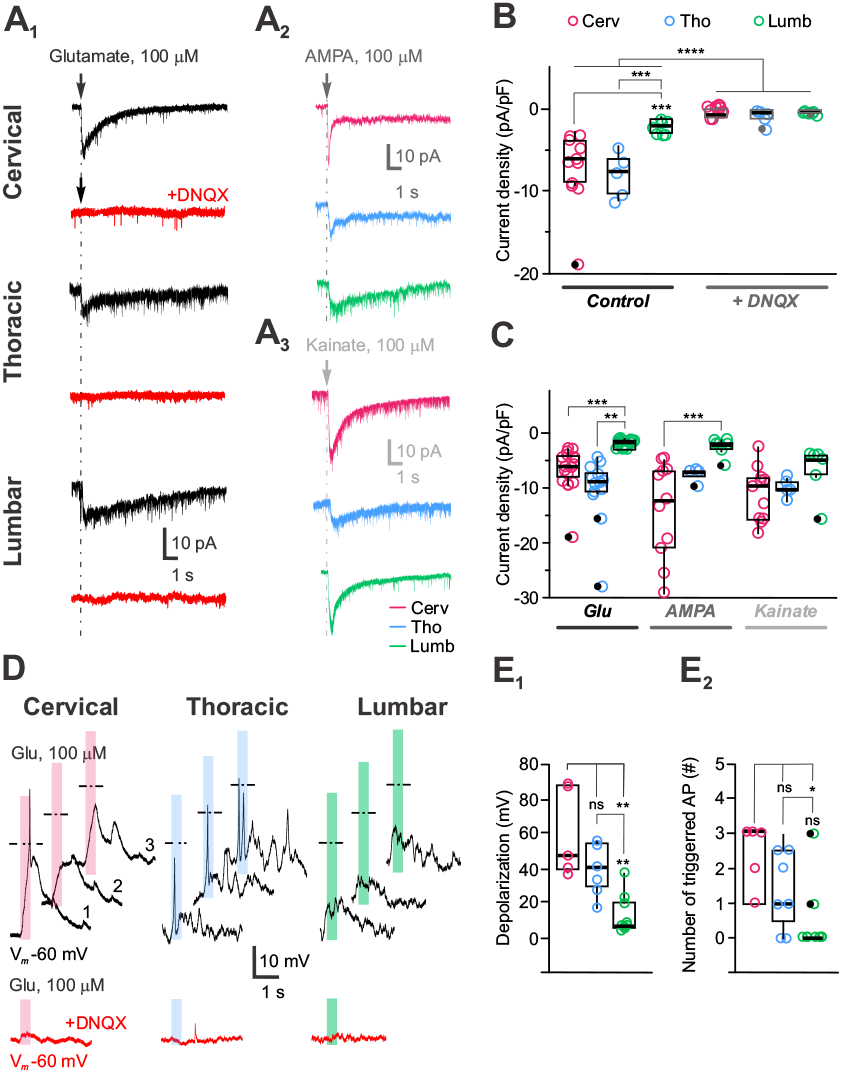
Slow kinetics-low amplitude AMPA/Kainate glutamatergic receptors modulate CSF-cN excitability. **A)** Representative whole-cell current traces recorded in CSF-cNs from the cervical, thoracic and lumbar segments (Top to Bottom) at V_h_ -80 mV in response to pressure application of glutamate (100 µM, 100 ms and 500 ms for the lumbar region; **A**_**1**_), AMPA (100 µM, 30 ms; **A**_**2**_) and Kainate (100 µM, 30 ms; **A**_**3**_). In **A**_**1**_, the representative currents recorded in the presence of DNQX (400 µM; red traces) upon Glutamate applications are illustrated for each segment below the control traces. Arrows and dashed lines in A_1_, A_2_ and A_3_ indicate tilme of agoist application. **B)** Summary box-and-whiskers plots of the averaged amplitude of the glutamate-mediated currents recorded at V_h_ -80 mV in control (Left) and in the presence of DNQX (Right) for the 3 segments of interest (-7±5, -8±3 and -2±1 pA.pF^-1^ in Control and 0.5±0.6, -0.8±1.0 and 0.2±0.2 pA.pF^-1^ in DNQX for the cervical (red, N=2, n=11), thoracic (blue, N=2, n=5) and lumbar (green, N=2, n=7) segments, respectively; ANOVA.lme: F=13.83, df=5 and 40, p(F)=7.362.10^−6^ and Tukey (EMM) post-hoc test to compare Control and DNQX currents within each region and Control vs. DNQX currents between the different regions; *** and **** for p<0.01 and <0.001, respectively). **C)** Summary box-and-whiskers plots of the averaged amplitude of the current densities recorded at V_h_ -80 in CSF-cNs from the cervical (red), thoracic (blue) and lumbar (green) in response to pressure application of glutamate (Glu), AMPA and Kainate (-7±4, -14±9 and -11±5 pA.pF^-1^; -10±6, -8±1 and -10±2pA.pF^-1^ and -2±1, -2±2 and -6±4 pA.pF^-1^ in cervical (red, N=3, n=16, 11 and 11), thoracic (blue, N=3, n=13, 4 and 5) and lumbar (green, N=3, n=17, 7 and 6) segments, respectively. Data and n are given Glu-, AMPA- and Kainate-mediated current, respectively; ANOVA.lme: F=7.907, df=8 and 81, p(F)=8.36.10^−8^ and Tukey (EMM) post-hoc test to compare Glu-, AMPA- and Kainate-mediated currents within each region and between the different regions; *, ** and *** for p<0.05, <0.01 and <0.001, respectively). **D)** Representative traces of the membrane potential changes recorded at V_m_ -60 mV in current-clamp mode in CSF-cNs from the cervical, thoracic and lumbar segments (Left to Right) in response to pressure application of glutamate (100 µM, 100 ms). Bottom traces in red are membrane potential changes recorded upon Glutamate application in the presence of DNQX. Shaded colored bars indicates the duration of the agonist application and dashed lines the 0 mV voltage. **E**_**1**_) Summary box-and-whiskers plots of the averaged depolarization levels of the membrane potential in CSF-cNs from the cervical (red, N=2, n=5), thoracic (blue, N=3, n=7) and lumbar (green, N=2, n=9) in response to pressure application of glutamate (Depolarization level: +60±26, +13±11 and +48±30 mV for C, T and L, respectively; Kruskal-Wallis rank sum test: 𝒳^2^=11.957, d=2, p(𝒳^2^)=2.533.10^−3^ and a post hoc Pairwise comparisons using Wilcoxon rank sum test with continuity correction: C vs. T, p=0.530; C vs. L and T vs. L, p=0.005). **E**_**2**_) Summary box-and-whiskers plots of the average number of AP triggered in CSF-cNs from the cervical (red, N=2, n=5), thoracic (blue, N=3, n=7) and lumbar (green, N=2, n=9) in response to pressure application of glutamate (AP number: 2.2±1.1, 1.4±1.3 and 0.4±1.0 for C, T and L, respectively; Kruskal-Wallis rank sum test: 𝒳^2^=7.5111, df=2, p(𝒳^2^)=0.02339 and a post hoc Pairwise comparisons using Wilcoxon rank sum test with continuity correction: C vs. T, p=0.302; C vs. L, p=0.0034 and T vs. L, p=0.117). In **B, D** and **E**, Single data points for all the recorded cells at each level are presented with colored opened circle (same color code as above).

Finally, we tested for functional nicotinic cholinergic receptors (nACh-Rs; see Corns and colleagues^39^) expression in CSF-cNs from all spinal segments. Acetylcholine application (ACh, 4 mM for 500 ms; aCSF and intracellular solution D, ***Supplementary Tables 1 and 2***) elicits current with an average amplitude of -9.5±8.0 pA and current density of -1.2±0.1 pA.pF^−1^ (N=28, n=65; ***Fig. 8A, B***) and was blocked by 94±19% (N=28, n=34; ***Fig. 8A***_***1***_, red traces, ***8B***) with D-tubocurarine (D-Tubo, 100-200 µM). T-CSF-cNs exhibit the smallest current amplitude (***Fig. 8B***). In current-clamp recordings at -60 mV (DC current injection of -10 to -20 pA; aCSF and intracellular solution A, ***Supplementary Tables 1 and 2***), ACh application depolarized the membrane potential by +23±15 mV (N=10, n=19; ***Fig. 8C, D***_***1***_), triggering APs except in T-CSF-cNs, likely due to their lower nACh-R-mediated current (***Fig. 8C, D***_***2***_). This depolarization was absent in the presence of D-Tubocurarine (***Fig. 8C***, Bottom) In contrast to findings in juvenile rats^9^, where CSF-cNs express functional P2X purinergic receptors, we were unable to replicate these results in mouse spinal CSF-cNs, as no ATP-γ-S-mediated current was detected.

**Figure 8.**
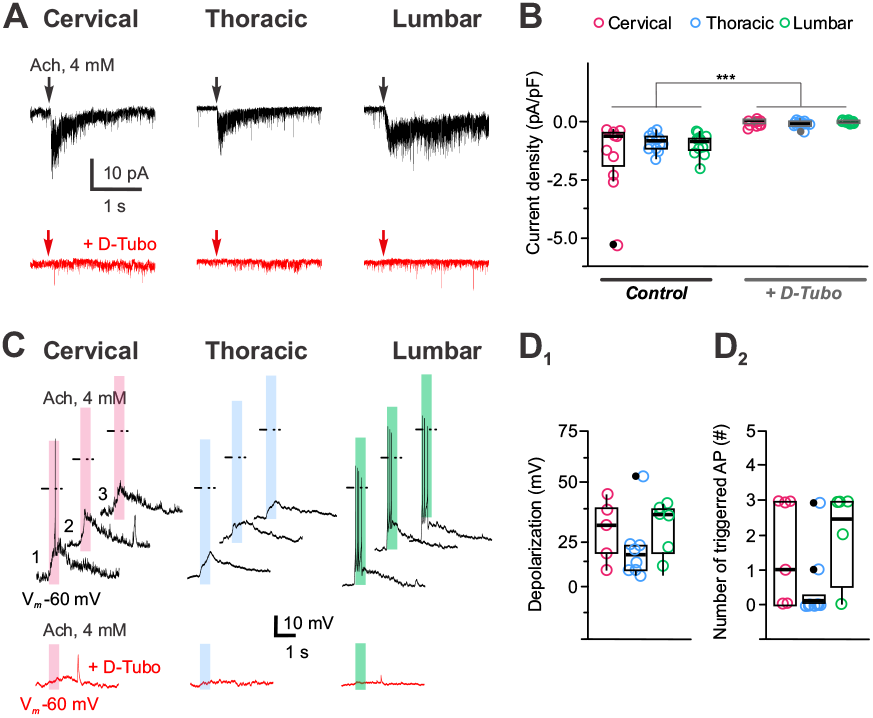
Nicotinic cholinergic receptors modulate CSF-cN excitability. **A)** Representative whole-cell current traces recorded in CSF-cNs from the cervical, thoracic and lumbar segments (Left to Right) at V_h_ -80 mV in response to pressure application of acetylcholine (ACh, 4 mM for 500 ms). The representative currents recorded in the presence of D-tubocurarine (D-Tubo, 100-200 µM; red traces) upon ACh applications are illustrated for each segment below the control traces. **B)** Summary box-and-whiskers plots of the averaged amplitude of the ACh-mediated currents recorded at V_h_ -80 mV in control (Left) and in the presence of D-tubocurarine (Right) for the 3 segments of interest (-1.4±1.5, -0.9±0.4 and -0.9±0.5 pA.pF^-1^ in Control and -0.0±0.1, -0.1±0.1 and 0.0±0.1 pA.pF^-1^ in D-Tubo for the cervical (red, N=8, n=11), thoracic (blue, N=13, n=12) and lumbar (green, N=7, n=11) segments, respectively; ANOVA.lme: F=9.192, df=5 and 62, p(F)=1.39.10^−6^ and Tukey (EMM) post-hoc test to compare Control and D-Tubo currents within each region and between the different regions; ** and *** for p<0.01 and <0.001, respectively). **C)** Representative traces of the membrane potential changes recorded at V_m_ -60 mV in current-clamp mode in CSF-cNs from the cervical, thoracic and lumbar segments (Left to Right) in response to pressure application of ACh (4 mM, 500 ms). Bottom traces in red are membrane potential changes recorded upon ACh application in the presence of D-Tubo. Shaded colored bars indicates the duration of the agonist application and dashed lines the 0 mV voltage. **D**_**1**_) Summary box-and-whiskers plots of the averaged depolarization levels of the membrane potential in CSF-cNs from the cervical (red, N=4, n=5), thoracic (blue, N=3, n=7) and lumbar (green, N=4, n=9) in response to pressure application of ACh (Depolarization level: 27±15, 2-18±15 and 27±15 mV for C, T and L, respectively; Kruskal-Wallis rank sum test: 𝒳^2^=1.3411, d=2, p(𝒳^2^)=0.5114 and a post hoc Pairwise comparisons using Wilcoxon rank sum test with continuity correction: C vs. T, p=1.0; C vs. L, p=0.62 and T vs. L, p=0.62). **D**_**2**_) Summary box-and-whiskers plots of the average number of AP triggered in CSF-cNs from the cervical (red, N=4, n=5), thoracic (blue, N=3, n=7) and lumbar (green, N=4, n=9) in response to pressure application of ACh (AP number: 1.4±1.5, 0.5±1.1 and 1.8±1.5 for C, T and L, respectively; Kruskal-Wallis rank sum test: 𝒳^2^=3.1265, df=2, p(𝒳^2^)=0.2095 and a post hoc Pairwise comparisons using Wilcoxon rank sum test with continuity correction: C vs. T, p=0.36; C vs. L, p=0.77 and T vs. L, p=0.33). In **B** and **D**, Single data points for all the recorded cells at each level are presented with colored opened circle (same color code as above).

Our data show that spinal CSF-cNs express functional GABA_A_, glycine, AMPA/Kainate glutamatergic, and nACh receptors. Activation of AMPA/Kainate and nACh receptors modulates CSF-cN excitability and can trigger AP firing.

### Voltage-gated calcium channels modulation by metabotropic receptors

Spinal CSF-cNs express functional ionotropic receptors for the major neurotransmitters (GABA, Glutamate and ACh) and in the *medulla*, we indicated that they also express their metabotropic subtypes capable of regulating Ca_V_ activity^19^. We investigated whether spinal CSF-cNs also express functional metabotropic receptors, and if these receptors modulate Ca_V_ activity postsynaptically. To test this, we elicited I_Ca_ at V_*h*_ -80 mV by applying a V_*Step*_ to +10 mV for 100 ms to record peak I_Ca_ (Ca_V_ modulation solution and intracellular solution B, ***Supplementary Tables 1 and 2***). Before each step, a 50 ms hyperpolarizing step to -120 mV was used to fully deactivate all Ca^2+^ channel subtypes. This protocol was repeated every 20 seconds under control conditions before pressure-applying agonists for a given metabotropic receptor.

We first assessed whether GABA_B_ receptor (GABA_B_-Rs) activation modulates Ca_V_ in spinal CSF-cNs (***Fig. 9***). In control conditions, the peak Ca^2+^ current (I_Ca_) had an average amplitude of -58±28 pA.pF^−1^ (N=8, n=31), which decreased to -40±21 pA.pF^−1^ (N=8, n=31) upon pressure application of Baclofen (Bcl, 100 µM, for 40 s) to selectively activate GABA_B_-Rs (***Fig. 9A, C***). The Bcl-mediated inhibition of I_Ca_ over time is shown in ***Figure 9B*** for CSF-cNs across the spinal regions. On average, Bcl inhibited I_Ca_ by 30±11% (N=8, n=31; ***Fig. 9D, Left***). This inhibition was reversible, with currents recovering to -51±26 pA.pF^−1^ (N=8, n=31), reaching 87±12% of control values. Although T-CSF-cNs had the largest I_Ca_ amplitudes, GABA_B_-R activation similarly inhibited currents across segments (***Fig. 9B-D***). In the presence of CGP54626A (2 µM, a GABA_B_-R antagonist), I_Ca_ amplitude was –55±29 pA.pF^−1^ (N=8, n=26), and Bcl application failed to inhibit I_Ca_ (average amplitude: -55±30 pA.pF^−1^, inhibition by 0±8%; N=8, n=26; ***Fig. 9D, Right***).

**Figure 9.**
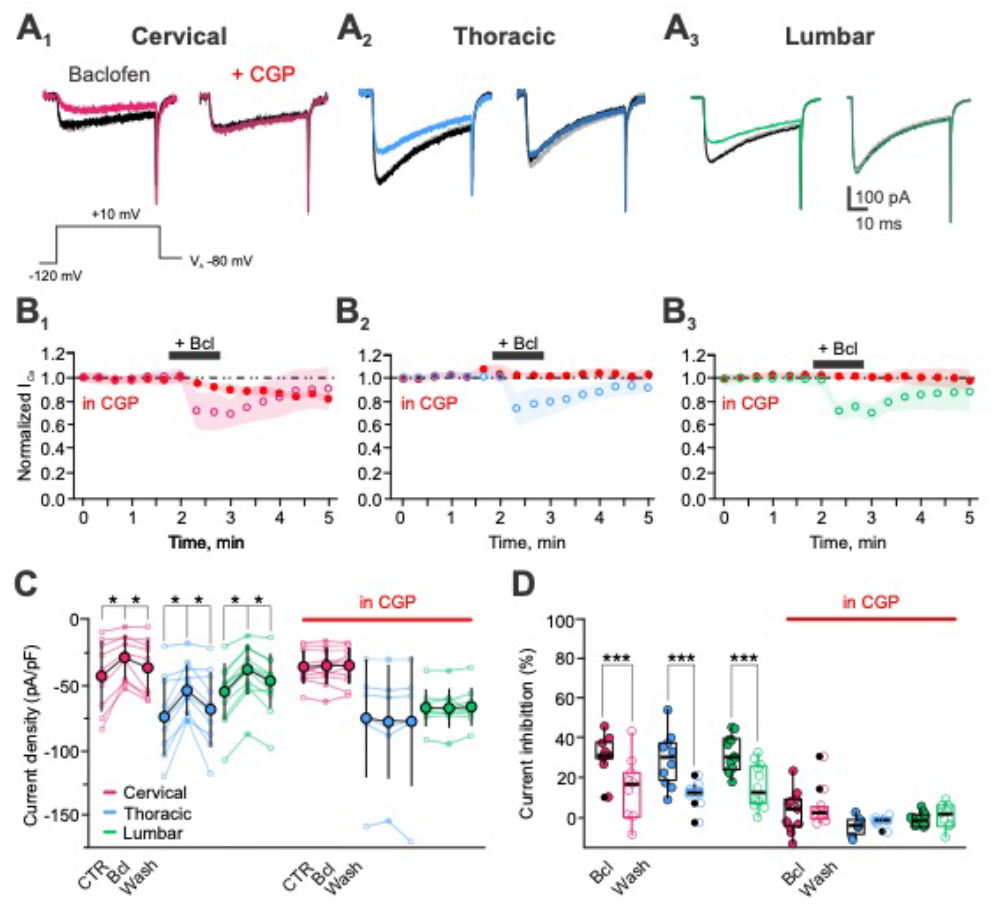
Metabotropic GABA receptors are functional in CSF-cNs and inhibit calcium channels. **A)**Representative I_Ca_ traces elicited at the peak with a voltage step to +10 mV from V_h_ -80 mV and recorded in CSF-cNs from the cervical (**A**_**1**_, red), thoracic (**A**_**2**_, blue) and lumbar segments (**A**_**3**_, green) in control (black traces), in response to pressure application of baclofen alone (Bcl; Left, colored traces) or in the presence of CGP (+CGP 2 μM; Right, colored traces) and after agonist Wash (grey traces). **B)** Averaged time courses of I_Ca_ peak amplitude recorded in CSF-cNs from the cervical (red; N=4, n=9), thoracic (blue; N=3, n=10) and lumbar (green; N=4, n=12) segments (from Left to Right). Bcl was applied at 100 μM by pressure for 40 s, as indicated by the black bar (+Bcl) and data were normalized to the baseline/control period before agonist application. Colored open circle: in Bcl alone; Red full circle: in the presence of CGP and SD: colored shaded area around the mean. **C)** Summary plots for the average peak I_Ca_ density before (CTR), during Bcl application and after agonist washout (Wash) in the absence (Left) or presence (Right, in CGP, red bar) of CGP. Filled colored circles: mean±SD; open colored circles: single data point; the color code represents the regions of interest: cervical (red), thoracic (blue) and lumbar (green). Bcl alone: -42±27, -28±17 and -36±22 pA.pF^-1^(N=2, n=9); -73±30, -53±20 and -67±29 pA.pF^-1^ (N=3, n=10) and -56±22, -39±19 and -48±21 pA.pF^-1^ (N=3, n=12) (Kruskal-Wallis rank sum test: 𝒳^2^=6.7051, df=2, p(𝒳^2^)=0.035 and a post hoc Pairwise comparisons using Wilcoxon rank sum test with continuity correction: CTR vs. Bcl, p=0.036; CTR vs. Wash, p=0.327 and Wash vs. Bcl, p=0.145). Bcl in CGP: -35±13, -34±14 and -34±14 pA.pF^-1^ (N=3, n=11); -74±45, -77±44 and -76±50 pA.pF^-1^ (N=3, n=16) and -66±14, -66±15 and -65±15 pA.pF^-1^ (N=3, n=9) (Kruskal-Wallis rank sum test: 𝒳^2^=0.026215, df=2, p(𝒳^2^)=0.987 and a post hoc Pairwise comparisons using Wilcoxon rank sum test with continuity correction: CTR vs. Bcl, CTR vs. Wash and Wash vs. Bcl, p=0.99, respectively). In the absence or presence of CGP, data are given in the order CTR, Bcl and Wash and for C, T and L, respectively. Note that amplitude for the I_Ca_ recorded in T-CSF-cNs are significantly larger (see **Fig. 5C**). **D**) Summary box-and-whiskers plots for the I_Ca_ inhibition induced before (Bcl) and after (Wash) agonist application and in the absence (Left) and presence (Right; in CGP, red bar) of CGP for the 3 regions of interest (C, red; T, blue and L, green). Bcl alone: 32±10 and 15±16% (N=2, n=9); 25±14 and 7±6% (N=3, n=10) and 32±9 and 16±11% (N=3, n=12) during and after Bcl application, respectively (Kruskal-Wallis rank sum test: 𝒳^2^=22.98, df=1, p(𝒳^2^)=1.637.10^−6^ and a post hoc Pairwise comparisons using Wilcoxon rank sum test with continuity correction: Bcl vs. Wash, p=3.6.10^−7^). Bcl in CGP: 3±11 and 3±13% (N=2, n=11); -5±5 and - 2±3 % (N=3, n=6) and 0±4 and 1±6% (N=3, n=6) (Kruskal-Wallis rank sum test: 𝒳^2^=0.90566, df=1, p(𝒳^2^)=0.3413 and a post hoc Pairwise comparisons using Wilcoxon rank sum test with continuity correction: Bcl vs. Wash, p=0.35). In the absence or presence of CGP, data are given in the order Bcl and Wash for C, T and L, respectively. Single data points for all the recorded cells at each level are presented with colored opened circle (same color code as above).

We next tested the effect of muscarinic cholinergic receptor (mACh-Rs) activation on I_Ca_ modulation. Using the same protocol as for GABA_B_-Rs, Oxotremorine-M (Oxo-M, 100 µM, selective mACh-R agonist) was pressure-applied while recording Ca^2+^ peak currents in C-, T-, and L-CSF-cNs (***Fig. 10***). The average Ca^2+^ peak current was -53±39 pA.pF^−1^ (N=9, n=36) in control and decreased to -31±25 pA.pF^−1^ (N=9, n=36) after Oxo-M application (***Fig. 10A-C***). On average, Oxo-M inhibited Ca^2+^ currents by 40±20% (N=9, n=31; ***Fig. 10D, Left***). This inhibition was reversible, with Ca^2+^ currents recovering to - 51±40 pA.pF^−1^ (N=9, n=36). The modulation was consistent across the CC axis (***Fig. 10B-D***). In the presence of atropine (Atr, 10 µM, a selective mACh-R antagonist), I_Ca_ amplitude was -38±30 pA.pF^−1^ (N=9, n=21), and Oxo-M) failed to inhibit Ca^2+^ currents (amplitiude of -36±28 pA.pF^-1^ and inhibition by 8±19%; N=9, n=21; ***Fig. 10D, Right***). We tested I_Ca_ modulation *via* metabotropic glutamatergic receptors (mGlu-Rs) by applying glutamate (100 µM, 40 s) but observed no reduction in current amplitude, indicating that Ca_V_ in CSF-cNs are not modulated by mGlu-R activation.

**Figure 10.**
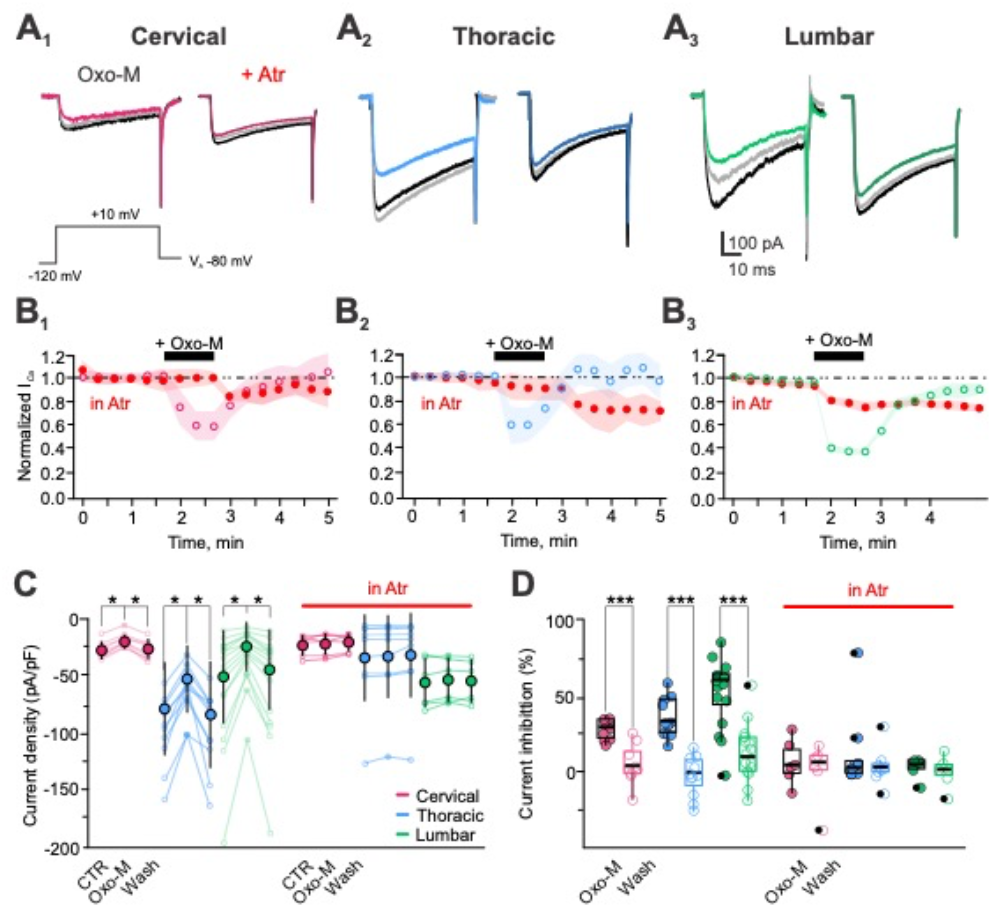
In CSF-cNs muscarinic activation reversibly inhibits calcium channels. **A**) Representative I_Ca_ traces elicited at the peak with a voltage step to +10 mV from V_h_ -80 mV and recorded in CSF-cNs from the cervical (**A**_**1**_, red), thoracic (**A**_**2**_, blue) and lumbar (**A**_**3**_, green) in control (black traces), in response to pressure application of Oxotremorine-M alone (Oxo-M; Left, colored traces) or in the presence of atropine (+Atr 10 μM; Right, colored traces) and after agonist Wash (grey traces). **B)** Averaged time courses of I_Ca_ peak amplitude recorded in CSF-cNs from the cervical (red; N=2, n=6), thoracic (blue; N=4, n=11) and lumbar (green; N=3, n=16) segments (from Left to Right). Oxo-M was applied at 100 μM by pressure for 30 s, as indicated by the black bar (+ Oxo-M) and data were normalized to the baseline/control period before agonist application. Colored open circle: in Oxo-M alone; Red full circle: in the presence of Atr (in Atr) and SD: colored shaded area around the mean. **C)** Summary plots for the average peak I_Ca_ density before (CTR), during Oxo-M application and after agonist washout (Wash) in the absence (Left) or presence (Right, in Atr, red bar) of Atr. Filled colored circles: mean±SD; open colored circles: single data point; the color code represents the regions of interest: cervical (red; N=2, n=9, 6), thoracic (blue; N=4, n=11, 9) and lumbar (green; N=3, n=16, 6) in Oxo-M alone: -27±7, -20±5 and -26±8 pA.pF^-1^; - 78±40, -52±29 and -83±46 pA.pF^-1^ and -50±40, -24±21 and -44±34 pA.pF^-1^ (ANOVA.lme: F=6.23, df=8 and 99, p(F)=1.654.10^−6^ and Tukey (EMM) post-hoc test to compare CTR, Oxo-M and Wash within a segment and between segments; *p<0.5). Oxo-M in Atr: -23±9, -22±9 and -21±8 pA.pF^-1^; -34±38, -33±36 and -32±37 pA.pF^-1^ and -56±22, -54±19 and -55±19 pA.pF^-1^ (ANOVA.lme: F=1.736, df=8 and 54, p(F)=0.1112 and Tukey (EMM) post-hoc test to compare CTR, Oxo-M and Wash within a segment and between segments). In the absence or presence of Atr, data are given in the order CTR, Oxo-M and Wash and for C, T and L, respectively. **D**) Summary box-and-whiskers plots for I_Ca_ inhibition induced before (Oxo-M) and after (Wash) agonist application and in the absence (Left) and presence (Right; in Atr, red bar) of Atr for the 3 regions of interest (C, red; T, blue and L, green). Oxo-M alone: 27±7 and 5±13% (N=2, n=9); 35±13 and -41±14% (N=3, n=11) and 51±23 and 6±32% (N=3, n=16) (ANOVA.lme: F=17.74, df=5 and 66, p(F)= 3.422.10^−9^ and Tukey (EMM) post-hoc test to compare inhibition in Oxo-M and during Wash within a segment and between segments; *,** and *** p<0.05, 0.01 and <0.001). Oxo-M in Atr: 7±15 and 1±20% (N=2, n=6); 13±25 and 4±12% (N=3, n=9) and 3±7 and 0±11% (N=3, n=6) (ANOVA.lme: F=0.5616, df=5 and 36, p(F)= 0.7286 and Tukey (EMM) post-hoc test to compare inhibition in Oxo-M and during Wash within a segment and between segments. In the absence or presence of Atr, data are given in the order Oxo-M and Wash for C, T and L, respectively. Single data points for all the recorded cells at each level are presented with colored opened circle (same color code as above).

In summary, our results show that all tested spinal CSF-cNs express functional GABA_B_- and mACh-Rs, whose activation inhibits post-synaptic Ca_V_. GABA_B_- and mACh-Rs activation induced a similar inhibition of I_Ca_ in all spinal CSF-cNs. Notably, we did not observe any modulation from mGlu-Rs, which have been shown to inhibit Ca^2+^ currents in other neuronal populations.

## DISCUSSION

CSF-cNs have been extensively studied in zebrafish larvae^5,27–29,34,35,40^, in lamprey^6,23,24^, and, to some extent, in mice^7,8,12,13,19,20,22,32^ and rat^9,11^. Our study provides an in-depth analysis of CSF-cN distribution and properties along the mouse SC from cervical to lumbar segments. We find that spinal CSF-cNs exhibit a conserved morphology, are densely distributed along the entire SC axis, and are primarily located ventrally. Functionally, they share intrinsic and chemosensory properties with mouse medullary CSF-cNs^12,22^. We show that spinal CSF-cNs express Na_V_ and K_V_ channels, LVA and HVA Ca^2+^ channels that are modulated by metabotropic GABAergic and muscarinic receptors but not by mGlu-Rs. They also express functional classical inhibitory and excitatory ionotropic synaptic receptors. Overall, spinal CSF-cNs represent a morphologically and functionally homogeneous sensory neuronal population.

### Spinal CSF-cNs form a dense morphologically homogenous neuronal network

Our study confirms and extends prior reports^7,8,13,20^, showing that in mice, CSF-cNs are distributed around the central canal (CC) along the entire rostrocaudal axis of the SC, exhibiting a characteristic and conserved morphology^5,7,9,11,32,33^. These neurons are either intra-ependymal, embedded within the ependymal layer, or subependymal, located below the CC, with a large dendrite extending into the CC lumen^7,9,11,33,33^. We show that CSF-cNs form a dense interconnected neuronal population, with approximately 10 to 20 cells per 10 µm of tissue depth across the entire CC axis. The SC length for a 6 weeks-old mouse and is ∼3.75 cm (3750 µm) and based on our confocal images the CSF-cN density is ∼14 cells/10 µm, then we estimate that ∼52,500 CSF-cNs would be present in the whole SC. Bjugn and Gundersen^41^ estimated that ∼6.4 million neurons are found in the whole SC grey matter and therefore spinal CSF-cNs would only represent 0.9% of the total SC neuronal population. Spinal CSF-cNs are predominantly found in the ventral region, comprising about 60% of the total. This ventral localization contrasts with the lateral distribution observed in the medulla^7^ (but see also Kútna and colleagues^32^) and suggest an anatomical reorganization along the medullo-spinal axis. We also confirm the presence of PKD2L1-expressing neurons (tdTomato^+^) in more distal ventral locations along the SC^13^. While we noted differences in the localization and density of cervical CSF-cNs, no functional difference was associated with these anatomical descriptions (see below).

Reports from studies in lamprey suggest that intra- and subependymal CSF-cNs represent two distinct subpopulations (type I and II)^6^. This property is also observed in embryonic and postnatal mice, where dorsally located CSF-cNs express Olig2 (CSF-cNs’) and latero-ventral ones express Nkx2.2 and Nkx6.1 (CSF-cNs”)^8,30^. A similar organization is observed in zebrafish larvae^29^. It has been suggested that ventral CSF-cNs may have a ‘more’ immature neuronal phenotype, which aligns with findings in older mice showing that CSF-cNs largely conserve an immature profile (expression of Nkx2.2, Nkx6.1, doublecortin and low levels of the Neuronal Nuclear protein)^7,8,12,29-31^. Nevertheless in older mice unlike in zebrafish or postnatal mice, medullo-spinal CSF-cNs subpopulations do not cluster specifically around the CC^7,13,31^, possibly due to developmental reorganization.

In zebrafish larvae, CSF-cNs have distinct projection paths and postsynaptic targets based on their dorso-ventral location, with ventral CSF-cNs showing longer projections, more branching, and a larger arborization area^29^. These neurons would have different postsynaptic targets^26,29,35,42^. Our study shows that CSF-cN axons project to the median fissure in the SC, forming long ascending fiber bundles in the lumbar to cervical segments. Nakamura and colleagues^14^ further demonstrated that CSF-cNs primarily project rostrally with a number of collaterals that is the highest in the dorsal part of cervical and the ventral lumbar segments. This higher density of CSF-cN projection in cervical and lumbar segments could be related to specific local spinal network and the associated function. Using selective viral retrograde tracing from the lumbar region, it was confirmed in mice that CSF-cNs project to rostral segments and form functional synaptic contacts with more rostral CSF-cNs^15^. This observation would further support the presence of two CSF-cN subpopulations in the SC. However, in mice CSF-cNs in ventral or dorsal regions could not be selectively labelled to demonstrate specific axonal projection paths *i*.*e*. postsynaptic targets relative to their localization around CC and rostro-caudal segments.

At the presynaptic level, it was shown that CSF-cNs project along the CC axis in an ascending manner to contact other CSF-cNs^15^ and monosynaptic retrograde tracing indicates that CSF-cNs are mainly contacted by GABAergic neurons although glutamatergic presynaptic partners have also been identified^15^. Nevertheless, the precise phenotype and localization within the spinal tissue of these presynaptic partners remains unknown and further investigation are needed to resolve this issue.

There might be phenotypical and anatomical evidence pointing towards the existence of two CSF-cN subpopulations in mice that would receive inputs from and project to specific presynaptic and postsynaptic partners, respectively, to date, no experimental data are available to support this organization and demonstrate its functional relevance in the different medullo-spinal segments. One might hypothesize that these identified subpopulations may differ by their integration in local networks (*i*.*e*. specific pre- and postsynaptic partners) to serve a modulatory function specific to a given medullo-spinal segment (see below). This hypothesis, that is investigated at the neuroanatomical level by several research groups in the field, is the first fundamental step to allow the characterization of CSF-cN functional connectivity and to allow the better understanding of CSF-cN function in the CNS.

### Along the spinal central canal, CSF-cNs exhibit similar cellular properties

CSF-cNs have been primarily studied at the behavioral level in zebrafish^5,27–29,34,35,40^ and to a lesser extent at the cellular one in mouse medullo-spinal tissues^12,19,20,22,33,39,43^. Across these models, CSF-cNs share common properties, including high input resistance in the Giga-ohm range and small membrane capacitance, in agreement with their small soma size and limited dendritic branching. Our data show that spinal CSF-cNs exhibit conserved intrinsic properties along the rostro-caudal axis, from the *medulla* to the lumbar SC, with no significant functional differences across spinal segments.

PKD2L1, a hallmark of medullo-spinal CSF-cNs^5,12,22,23,28^, is confirmed in our study as being expressed and functional along the SC. This non-selective cationic channel is modulated by extracellular pH and can trigger APs, influencing CSF-cN excitability^22^. We found that PKD2L1 is activated by alkaline pH and inhibited by acidic pH, while spinal CSF-cNs also respond to acidic pH via ASICs activation. Both mechanisms affect CSF-cN excitability, supporting a conserved sensory function along the CC. Although chemosensitivity has been demonstrated using non-physiological stimuli, further research is needed to identify the receptors that respond to chemical clues present in the CSF.

We observed that T- and C-CSF-cNs exhibit a depolarized RMP of approximately -50 mV, consistent with previous findings for medullar CSF-cNs^12,22^. In contrast, L-CSF-cNs display a more hyperpolarized RMP of around -65 mV, a result recently reported using the less invasive cell-attached recording technique^33^. This suggests that L-CSF-cNs may have different ionic conductances or reduced PKD2L1 activity, contributing to their hyperpolarized membrane potential. However, our study shows no differences in the expression of major K_V_ channels or PKD2L1 activity.

In postnatal mice^8^ and juvenile rat^9^, CSF-cNs were shown to form two major neuronal subpopulations present in different region around the CC, exhibiting different state of maturity and firing patterns (see above). In older mice, we have shown that two medullar CSF-cN population can be distinguished based on their maturity state and firing pattern^12,22^ an observation confirmed for spinal CSF-cNs in this study. However, in older mice and in contrast to the data obtained in zebrafish or postnatal mice^8^, medullo-spinal CSF-cNs are not regrouped in specific clusters but rather distributed around the CC. One can argue that during development and maturation CSF-cNs may redistribute around the CC or might undergo developmental process to change their phenotypical and functional properties. Dedicated developmental studies are required to address this specific point. We also observed that C- and L-CSF-cNs exhibit mainly a single spike firing pattern while thoracic neurons fire mostly tonically. This result suggests different properties and ionic conductances expression for CSF-cNs along the rostro-caudal axis. This might also depend on the specific network CSF-cNs are inserted in as well as to the exposure to given bioactive signals. Further, CSF-cNs are present in the so-called spinal niche in contact with the CSF and appear to interact with ependymal cells^10^. One could suggest that they might change functional properties as a function of the physio-pathological state as recently demonstrated following SC injury^10^. The observed difference could also be explained by the properties of the networks CSF-cNs are inserted in as cervical and lumbar segments contain motor networks mostly exhibiting phasic activities (locomotor output) while in thoracic regions autonomous centers would have a more sustained activity. However, one has to remain cautious in concluding about AP discharge activity when using whole-cell patch-clamp technique especially in neurons with small somatas. An answer to this point could be addressed *in vivo* using imaging approaches (calcium and voltage sensitive probes) or electrophysiological recordings.

Marichal and colleagues^9^ demonstrated that in juvenile rats CSF-cNs express functional TTX-sensitive Na^+^ and K^+^ voltage-dependent channels, including delayed-rectifier (K_DR_) and A-type (K_V_ 1) channels. Our findings align with these reports, showing that spinal CSF-cNs express Na_V_ channels at similar densities, mediating only transient and TTX-sensitive currents, as we did not observe any persistent or TTX-resistant Na^+^ currents. Regarding K_V_, all CSF-cNs exhibit large K^+^ currents, characterized by a fast, transient phase followed by a persistent one. In agreement with previous reports^9^, our pharmacological analysis reveals that spinal CSF-cNs express 4-AP-sensitive fast and transient currents along with TEA-sensitive persistent currents. This indicates that, as in juvenile rats^9^, mouse spinal CSF-cNs express K_DR_ and K_v_ 1 channels at similar densities. These Na_V_ ad K_V_ channels likely underlie AP generation in spinal CSF-cNs. However, unlike Marichal and colleagues^9^, we did not observe differences in current amplitude or density based on the firing pattern (single spike *vs*. tonic) of CSF-cNs. It is possible that modulation of these channels by intracellular signaling pathways, potentially activated by agents circulating in the CSF, could differentially modulate CSF-cN excitability. Such modulation might directly affect AP properties or indirectly influence them through PKD2L1, which could subsequently trigger AP firing.

In juvenile rats^9^ and more recently in the mouse thoracic SC^20^, CSF-cNs were shown to express functional Ca_V_ channels, including the T-type LVA subtype (Ca_V_ 3). We previously reported the absence of Ca_V_ 3 channels in mouse medullary CSF-cNs, where N-type channels (Ca_V_ 2.2, HVA) were highly expressed^19^. However, in mouse spinal CSF-cNs, we now demonstrate HVA expression along with LVA channel subtypes as indicated by the characteristic ‘current shoulder’ observed when neurons were held at -80 mV, but not -60 mV, prior to Ca_V_ activation. The presence of T-type Ca^2+^ channel might contribute to CSF-cN excitability as well as Ca^2+^ signaling^33^, particularly through rebound burst firing after inhibitory synaptic inputs that hyperpolarize the cells, enabling T-type channel activation. We also found that thoracic CSF-cNs had the highest expression level of Ca_V_ channels, as reflected in the larger amplitude of I_Ca_ densities recorded in this region. This suggests that Ca^2+^ signaling is differentially regulated along the SC axis, with higher Ca_V_ density in thoracic segments.

We investigated the modulation of CSF-cN activity and Ca^2+^ signaling by examining Ca_V_ modulation via G protein-coupled receptors (GPCRs). Consistent with previous findings in the brainstem^19^, we found that Ca_V_ channels in all spinal CSF-cNs are reversibly inhibited following GABA_B_-R activation and also indicate a modulation by mACh-Rs. Both GABAergic and cholinergic GPCR activations led to similar levels of Ca_V_ inhibition, likely mediated through the classical Gβγ pathway^44^. Interestingly, we did not observe any Ca_V_ inhibition via glutamatergic GPCRs, suggesting that either these receptors are not expressed in CSF-cNs or they do not modulate Ca_V_ in this population. As reported for medullary CSF-cNs^19^ and more recently in lumbar segments^33^, we found no evidence for functional GIRK channels in spinal CSF-cNs, as GABA_B_-R activation failed to induce outward currents in all recorded neurons.

Here, we confirm our previous findings in medullary CSF-cNs and show that PKD2L1 is expressed in CSF-cNs along the SC axis, exhibiting spontaneous activity and contributing to CSF-cN excitability^22^. Supporting their chemosensory role^22–24,27,28^, we report that spinal CSF-cN excitability is modulated by extracellular acidification through activation of ASICs or by alkalinization *via* increased PKD2L1 channel open probability. CSF-cNs are capable of differentiating pH changes by responding to both increases and decreases in pH through specific ionic channels. It is likely that bioactive compounds in the CSF activate intracellular pathways, modulating CSF-cN activity either directly or indirectly through PKD2L1 or other receptors^10,36^, though these compounds (such as neurotransmitters, metabolites, or neuropeptides) remain largely unidentified in mice. In zebrafish larvae, PKD2L1 integrates mechanical signals by detecting CSF flow and SC bending, interacting with the Reissner fiber^27,28^. A similar mechanosensitivity was suggested in mouse thoracic CSF-cNs, although strong mechanical stimuli were required to modulate their excitability^20^. We were not able to confirm this result, since in our hands pressure application of agonists or aCSF failed to induce changes in PKD2L1 activity. Identifying the precise sensory stimuli to which CSF-cNs respond remains a critical open question in mice and is essential for understanding their function.

### CSF-cNs express classical synaptic receptors to be player of intraspinal networks

CSF-cNs might integrate circulating signals to sense the CNS homeostatic state. On the other hand, they appear to be integrated along the CC within intramedullary and spinal networks, which are involved in regulating CNS activity and body physiology. Recent studies^14,15^ have begun to explore CSF-cN connectivity, but limited information is still available regarding their presynaptic partners. In mice, medullary CSF-cNs express GABA_A_, glycine, and glutamatergic receptors and primarily exhibit spontaneous GABAergic and glycinergic synaptic activity at low frequencies (∼1 Hz)^19^ and both GABAergic and glutamatergic synaptic currents were evoked in medullary CSF-cNs^19^. However, the identity and localization of these presynaptic partners remain unknown. Our study shows that along the spinal CSF-cNs also express functional GABA_A_, GABA_B_, and glycine receptors, with large inhibitory currents elicited by exogenous agonists, suggesting robust inhibitory control. In the lumbar region, CSF-cNs were found to form two subpopulations, one depolarized and the other hyperpolarized by GABA_A_ receptor activation^33^. Although this has not yet been confirmed in thoracic and cervical regions, it is plausible that similar subpopulations exist. Furthermore, we show that spinal CSF-cNs express AMPA/kainate-type glutamatergic receptors, whose exogenous activation induces small, slow-kinetic currents, consistent with reports from the medulla^19^. This contrasts with the large, fast AMPA currents seen in other neurons or those electrically evoked in medullary CSF-cNs. Additionally, CSF-cNs along the spinal CC axis express muscarinic and nicotinic cholinergic receptors. Despite the small amplitude of these glutamatergic and cholinergic ionotropic currents, due to the high CSF-cN input resistance, receptor activation can induce significant depolarization and even trigger APs. Thus, activation of glutamatergic and cholinergic ionotropic receptors in CSF-cNs likely modulates their excitability. Within the spinal tissue numerous GABAergic, glutamatergic and cholinergic interneurons are present and shape spinal activity and outputs (somatic and autonomous motor systems). Recent reports^14,15^ and our data indicate that CSF-cNs express the major synaptic receptors and appear receiving inputs from or project to spinal interneurons. Therefore, CSF-cNs might be key player in the SC circuitry in the mammals by bidirectionally interacting with different neuronal populations to regulate CNS activity.

Spontaneous synaptic activity has been observed in medullar CSF-cNs, but not in spinal CSF-cNs (our study), raising questions about the source of neurotransmitters and their release conditions in activating spinal CSF-cNs. Neurotransmitters might be released via synaptic contacts with glutamatergic, GABAergic, and cholinergic neurons within the SC, nevertheless, except for recurrent connectivity among CSF-cNs as source for GABA^14,15^, functional contacts with CSF-cNs remain to be definitively demonstrated. Another potential origin could be paracrine release from neurons in the surrounding parenchyma, but this seems unlikely, as no baseline current changes were observed upon exposure to selective antagonists. However, one cannot rule out a loss or ‘dilution’ of such paracrine transmission in *in vitro* models where slice preparation is perfused with aCSF. Additionally, neurotransmitters have been shown to be present in the CSF and might activate CSF-cNs via their protrusions as suggested by a recent study indicating that CSF-cNs can be activated by κ-opiods released by neighboring cells^10^. Such a route of activation remains to be demonstrated. Nevertheless, there are growing anatomical evidence indicating that CSF-cNs appear integrated within specific spinal network and one major challenge in the coming years will be to further characterize the presynaptic partners of CSF-cNs and understand how they are activated. Gerstmann and colleagues^15^ used monosynaptic retrograde neuronal tracing to show that L-CSF-cNs are primarily contacted by GABAergic neurons, with some glutamatergic input as well. Extending this type of characterization to the entire SC and testing functional connectivity *in vivo* will be crucial.

### Concluding remarks

Altogether, the data collected in this study contribute to the better understanding CSF-cNs functions and suggest they represent a homogenous population present along the medullo-spinal axis that form an interconnected neuronal network to act as a coordinated sensory system. Nevertheless, the signals circulating in the CSF or present in the parenchyma that they respond to in physiological or pathological states remain to be fully identified. In physiological and pathological conditions, CSF-cNs would integrate sensory signals and in turn modulate, along the medullo-spinal axis, specific networks regulating or controlling either motor^14,15,26,34,35^, autonomous or even regenerative functions^10^.

As such, we propose that CS-cNs represent a novel actor of the interoceptive system capable of informing the CNS about its state and at the same time modulating specific physio-pathological outputs according to the segment there are located in.

The challenge in the coming years to demonstrate the role of CSF-cNs as interoceptors will be to characterize their connectivity, since, as Kolmer suggested it “*In many of these cases, precise anatomical knowledge was a prerequisite for physiological research*”^45^.

## MATERIAL AND METHODS

### Animal ethics / Ethical approval

All experiments were conducted in conformity with the rules set by the *EC Council Directive* (2010/63/UE) and the French “*Direction Départementale de la Protection des Populations des Bouches-du-Rhône* (DDPP13)” (Project License Nr: APAFIS 44331,2023071917567777 & 33336, 202110181902 2439. WN and License for the Use of Transgenic Animal Models Nr: DUO-5214). Protocols used agree with the rules set by the *Comité d’Ethique de Marseille* (CE71), our local Committee for Animal Care and Research. All animals were housed at constant temperature (21°C), in an enriched environment, under a standard 12h light-12h dark cycle, with food (pellet AO4, UAR, Villemoisson-sur-Orge, France) and water provided *ad libitum*. Every precaution was taken to reduce to the minimal the number of animals used and minimize animal stress during housing and prior to experiments.

### Animal models

We used wild type C57 Black6J (Charles River), PKD2L1-Cre (Pkd2l1^tm1(cre)^; MGI ID: 6451758; a generous gift Emily Leman), and Choline Acetyl Transferase-Cre (ChAT-Cre; Chat^tm2(cre)Lowl^; The Jackson Laboratory, MGI ID: 5475195; RRID:IMSR_JAX:006410) mice. PKD2L1- and ChAT-Cre animals were cross-breed with *flex*-tdTomato mice (Gt(ROSA)26^Sortm14 (CAG-tdTomato)Hze^, The Jackson Laboratory, MGI ID: 3809524; RRID:IMSR_JAX:007914) to generate PKD2L1-Cre (PKD-tdTomato)- and ChAT (ChAT-tdTomato)-Cre::*flex*-tdTomato mice and to selectively express the tdTomato fluorescent protein in the neuronal population of interest. Animal of either sex were used for histology as well as for electrophysiological recordings (3-6 Weeks old mice).

### AAV injections

3–5-week-old mice were deeply anesthetized with isoflurane and placed in a stereotaxic frame. Meloxicam (5 mg.Kg^-1^; Metacam, Boehringer Ingelheim),), a non-steroïdian anti-inflammatory compound, was administered intraperitoneally at least 30 minutes before surgery. Before making a skin incision, lidocaine (5 mg.Kg^-1^; Lurocaïne, Vetoquinol) was administered subcutaneously under the zone to be incised. Craniotomy was performed using a motorized drill (78001 Microdrill, RWD life sciences) using drill bits diameters of 1 mm (WPI) in the following coordinates (AP 0.0, ML 1.0 mm; DV 1.6 mm) and 700 nL of AAV1-hSyn-EGFP viral particles at 2×10^13^ GC.mL^-1^ were delivered using a Nanoject-III system (Drummond Scientifc Company) at a rate of 4 nL.s^-1^.

### Histology and imaging

#### Tissue collection for imaging and microscopy

Animals were injected 30 minutes prior to the procedure with Meloxicam (5 mg/Kg 30 min prior to procedure). Subsequently mice were anesthetized with intraperitoneal administration of 100 mg/Kg ketamine (Virbac) and 10 mg/Kg xylazine (Rompun, Bayer). Before skin incisions, the local antalgic lidocaine (5 mg/Kg; Lurocaïne, Vetoquinol) was injected subcutaneously. Afterwards, animals were transcardially perfused with 20 mL of ∼37°C PBS 0.1M followed by 20 mL of ice-cold 4% paraformaldehyde (EMS, 15714S) in PBS 0.1M. SC were removed by laminectomy and the meninges were gently separated prior to overnight fixation at 4°C in the same fixative solution. For clearing experiments, brain plus SC were dissected and post-fixed overnight at 4°C in the same fixative solution under gentle shaking.

#### Routine histology and confocal microscopy

SC were cut in three segments corresponding to cervical, thoracic and lumbar regions. Afterwards, each one of the segments was included in a low melting point agarose block (4% in PBS). These blocks were glued to a vibratome (HM 650V, Microm, Thermo Scientific) stage, and coronal slices (50 μm) of SC were collected as floating sections. After incubation in 1.5μg/mL of DAPI (D9542, Sigma-Aldrich) in 0.1M PBS (15 min, RT), sections were washed one last time in PBS and mounted in Vectashield (H-1900-10, Vector laboratories). Slides were kept at 4°C until use. 8-bits z-stacks images were acquired on a Zeiss LSM-700 Confocal scanning microscope equipped with ZEN 2009 software (Zeiss) using an EC Plan-Neofluar 10x, 0.30 NA objective, zoom=0.7x, or a Plan-Apochromat 20x, 0.8 NA objective, zoom=1x or 1.5x. Optical thickness was 4.45 μm for 10x acquisitions and 0.75 μm for 20x acquisitions.

#### Clearing and light sheet microscopy

For SC clearing, the vDISCO protocol^37,38^ was followed with minimal modifications. Since SCs are highly myelinated tissues, to facilitate the access of nanobodies to CSF-cNs, a detergent-based delipidation step derived from the SHANEL^46^ protocol was pre-applied before boosting the endogenous tdTomato signal. Briefly, brain plus SCs were incubated in a solution containing 10% wt/vol 3-[(3-cholamidopropyl)dimethylammonio]-1-propanesulfonate (CHAPS, 1479.1 Carl Roth), 25% vol/vol N-Methyldiethanolamine (N-MDEA, 471828 Sigma-Aldrich), 0.05% Sodium Azide in Type 1 diH20 during 24 h at 37°C. After washing in 0.1M PBS (3 times; 2h/ea), samples were transferred to the permeabilization solution containing 0.5% vol/vol Triton X-100 (T8787, Sigma-Aldrich), 5mM Methyl-β-Cyclodextrin (332615, Sigma-Aldrich), 0.3M Glycine (G7126, Sigma-Aldrich), 1.5% vol/vol Normal Goat Serum (S-1000, Vector laboratories), 0.05% wt/vol Sodium Azide in 0.1M PBS, and incubated 48h at 37 °C. Subsequently, samples were transferred to permeabilization solution containing 1:600 vol/vol FluoTag anti-RFP Alexa647 (N0404-AF647-L, NanoTag Biotechnologies; RRID:IMSR_JAX:007914) and incubated during 5 days at 37 °C. Afterwards, samples were washed 3 times for 2h/ea, and once o/n in washing solution containing 0.5% vol/vol Triton X-100 (T8787, Sigma-Aldrich), 1.5% vol/vol Normal Goat Serum (S-1000, Vector laboratories) in 0.1M PBS at RT. Then, samples were washed 4 times (2h/ea) in 0.1M PBS at RT and the clearing process was immediately started. To preserve the SC in a linear disposition, samples were gently stretched and pinned with 0.15mm-diameter stainless steel entomology pins to a thin 4% wt/vol low melting agarose block before clearing. For clearing, samples were incubated at RT in the following gradient of Tetrahydrofuran (THF, 186562, Sigma-Aldrich) in Type 1 H20, 1h/ea step: 50% vol/vol THF, 70% vol/vol THF, 80% vol/vol THF, 100% vol/vol THF, and o/n in 100% THF. After dehydration, samples were incubated in Dichlorometane (DCM, 270997, Sigma-Aldrich) 1h at RT, and finally in Benzyl Alcohol-Benzyl Benzoate 1:2 vol/vol (BABB, 305197 and B6630 respectively, Sigma-Aldrich) at RT until transparency. All incubations were performed under gentle shaking on an orbital shaker. Prior to imaging, samples were transferred to Ethyl cinnamate (ECI, W243000, Sigma-Aldrich) since it was the medium used for imaging.

#### Light-Sheet microscopy imaging

16-bits image stacks were acquired using an Ultramicroscope Blaze (Miltenyi Biotec) equipped with the following lasers and filter sets [laser (emission/bandwidth)]: 445(480/40), 488(525/50), 515(540/30), 561(595/40), 639(680/30). For acquisition of the autofluorescence signal (to get the whole morphology of the brain plus SC) the 445(480/40) configuration was used; for acquisition of the boosted tdTomato signal, 639(680/30) configuration was used. The following objectives were used depending on the acquisition needs: 1.1x objective, 0.1 NA, ≤ 17mm WD; 4x Objective, 0.35 NA, ≤ 17mm WD; 12x Objective, 0.35 NA, ≤ 11mm WD. When Tile scans were obtained, a minimum of 20% overlapping was used to ensure a proper stitching of the tiles. Exposure time was 100-200 ms and laser power was adjusted depending on the intensity of the fluorescent signal (to prevent saturation) and on the magnification used. The light-sheet width was kept at 80% for low magnification imaging and at 20% for high magnification imaging. The z-Stacks shown in figure 1A have a voxel size of 8.21 × 8.21 × 7.09 μm (x, y, z respectively). The z-Stacks shown in figure 1 B, 1C and 1D have a voxel size of 1.625 × 1.625 × 2 μm (x, y, z respectively). The z-Stack shown in ***Supplementary Fig*.*2C, D*** has a voxel size of 0.65 × 0.65 × 2 μm (x, y, z respectively).

### Image processing

Image analysis and rendering was performed on a workstation equipped with an 8 core Intel Xeon Silver 4215R processor (128Gb RAM) and a NVIDIA GeForce RTX 3090 (24 Gb) graphics card. FIJI^47^ and Chimera X^48,49^ were used for image processing, visualization, and rendering. Light-sheet image tiles were stitched with BigStitcher^50^ plugin and further processed in FIJI. For Image rendering, a rolling ball background subtraction^51^ was used over both channels with a radius size according to the particles of interest (Width of the SC in 445(480/40) and diameter of CSF-cNs axon bundles in 639(680/30)) in the different images. Then, the channel containing the specific signal (AF647) was subsequently treated with Contrast Limited Adaptive Histogram Equalization (CLAHE) in FIJI using a block-size larger (in pixels) than the objects to be preserved in order to compensate fluorescence discrepancies between CSF-cNs somas and thin processes. The background signal corresponding to the surface of the SC was removed by thresholding the autofluorescence signal and creating a reduced selection (in pixels) over the tdTomato (AF647) signal. The function “Clear Outside” was then used in FIJI over this selection to remove surface background fluorescence ensuring that objects in the specific channel (AF 647) were preserved. This strategy did not compromise the objects of interest since CSF-cNs and their axon bundles are distant from the surfaces of the SC. Finally, for image rendering, an “Unsharp Mask” filter was used with a radius of 0.9 pixels and a Mask Weight of 0.7. Rendering was performed in Chimera X by loading the previously generated .tiff stacks and adjusting graphical parameters. “Maximum Intensity Projection” and “3d projection mode” were used when displaying volumes. For low-magnifications large field of view images in ***Figure 2***, individual z-stacks were acquired as positions and stitched through Grid Collection Stitching plugin in FIJI^52^ to avoid edge artifacts derived from Tile acquisition in ZEN software. For high-magnification images in ***Figure 2***, Images were processed with CLAHE in order to avoid fluorescence discrepancies between the somas and the thin axonal processes of CSF-cNs since linear adjustment will inevitably result in saturation at the somatic level.

### Acute spinal cord slice preparation

Coronal SC slices (from lumbar to cervical segments) were prepared as previously described^19^. Briefly, wildtype or transgenic mice were injected with Meloxicam (5 m/Kg; 30 min prior to procedure). Subsequently, they were anaesthetized with intraperitoneal injection of ketamine–xylazine mixture (100 and 15 mg/kg, respectively) and injected on the site of surgery with Lurocaine (5 m/kg), a local antalgic. Next, animals were transcardially perfused with an ice-cold (0-4°C), oxygenated (95% O_2_/5% CO_2_) and low calcium/high magnesium slicing solution containing (in mM): NaCl 75, NaHCO_3_ 33, NaH_2_PO_4_ 1.25, KCl 3, CaCl_2_ 0.5, MgSO_4_ 7, glucose 15, sucrose 58, ascorbic acid 2, Na-pyruvate 2, myo-Inositol 3 (pH 7.3-7.4 and osmolarity of 300-310 mosmole.kg^-1^). Following laminectomy and SC dissection, lumbar to cervical SC slices (250 µm thick) were cut with a vibratome (Leica VT1000S) in ice-cold cutting solution saturated with 95% O_2_-5% CO_2_. Slices were subsequently transferred to a recovery chamber and incubated at 35° C for 15-20 minutes in oxygenated artificial CSF (aCSF) containing (in mM): NaCl 115, NaHCO_3_ 26, NaH_2_PO_4_ 1.25, KCl 3, CaCl_2_ 2, MgSO_4_ 2, glucose 15, ascorbic acid 2, Na-pyruvate 2, myo-Inositol 3 (pH 7.3-7.4 and osmolarity of 300-310 mosmole.kg-1) and then kept at room temperature to recover for 1 h. After recovery, slices were transferred one by one to the recording chamber.

### Electrophysiological recordings

#### CSF-cN visualization and recording of their intrinsic properties

For recordings, lumbar, thoracic, or cervical SC acute slices were transferred in a recording chamber mounted on an upright microscope (Zeiss Axioskop 1FS or Scientifica SliceScope Pro 1000) equipped with infra-red differential interference contrast illumination (IR-DIC or IR-Oblic) and a p1 or p300 Ultra precisExcite LED epifluorescence system (CooLED). CSF-cNs around the CC were visualized using a computer controlled digital camera (HQ2 CoolSnap, Photometrics, SciCam, Scientifica or XX) under epifluorescence illumination (exc. 520 nm/em. 610 nm, tdTomato fluorescence) and/or IR illumination with a 40x or 60x objective. Whole-cell patchclamp recordings were performed at room temperature (∼20°C) in current- and/or voltage-clamp mode using a Axopatch 200A or Multiclamp 700B amplifiers (Molecular Devices Inc.). Patch pipettes (3-7 MΩ) were pulled from borosilicate glass capillaries (OD: 1,5 mm, ID: 0,86 mm; Harvard Apparatus) using the vertical PC-100 puller (Narishige International Ltd) and filled with an internal solution (see detail in ***Supplementary Table 1***). Signals were low-passed filtered at 2-10 kHz and digitized between 10-50 kHz using a Digidata 1322A, 1440A, and 1500B interface driven by pClamp 9, 10 or 11 software (Molecular Devices). Series resistance (10-20 MΩ) was monitored throughout the experiment and unstable recording were discontinued (increase by more than 25% from its initial value). For voltage-clamp recordings, the series resistance was compensated by 70-80%. The liquid junction potential was left uncorrected. SC slices were perfused with oxygenated, and pH controlled (95% O_2_/5% CO_2_) artificial CSF (aCSF, 2.5 mL/min) and the composition was adapted according to the experiments conducted (see ***Supplementary Tables 1 and 2***).

The recording of CSF-cNs was confirmed based on the characteristic morphology observed (small round soma close or within the ependymal layer and a large dendrite ending in the CC with a round protrusion) from the cytosolic tdTomato as well as from Alexa488 or 594 Hydrazide (added to the intracellular solution, 10 µM; Invitrogen) fluorescence. The presence of spontaneous PKD2L1 channel activity was monitored as a further control. Resting membrane potential (RMP) was determined under current-clamp mode (I=0) and CSF-cN spontaneous or induced (step of 200-500 ms, -20 pA with increments of 10 pA, 10-12 sweeps) AP discharge were recorded in current-clamp mode either at RMP or a membrane potential set at -60 mV (current injection of - 10 to -15 pA). The intrinsic properties (membrane resistance, r_*m*_; time constant, τ and membrane capacitance, c_*m*_) were determined from the current response elicited using a 10 ms voltage step to -10 mV from a holding potential set at -70 mV in voltage-clamp mode. Note that to account for potential difference in ionic channel and receptor expression as a function cell size, all current amplitudes are expressed as current density (current/*C*_*m*_).

The characterization of CSF-cN modulation by pH was conducted either in voltage-clamp mode (V_*h*_ -80 mV) or current-clamp mode at RMP using standard aCSF complemented with antagonists against GABA_A_, glycine, and glutamate receptors and a K-based intracellular solution (in mM: KCl 130; NaCl 10; CaCl_2_ 1; MgCl_2_ 1, HEPES 10, phosphocreatine 10; Mg-ATP 4; pH 7.35adjusted with KOH and osmolarity of 290 mosmole.kg^-1^). Acidic (pH 5 (mM): NaCl 139; KCl 3; MgCl_2_ 2; CaCl_2_ 2; glucose 15; HEPES, 10, Na-Citrate 16; pH adjusted with HCl 1N and 310 mosmole.Kg^-1^) or alkaline (pH 9 (mM): NaCl 145; KCl 3; MgCl_2_ 2; CaCl_2_ 2; glucose 15; HEPES 10, TAPS 10; pH adjusted with NaOH 1M and 310 mosmole.Kg^-1^) solutions were applied by pressure for 10 (pH5) or 30 s (pH 9) using a glass patch pipette positioned close to the recorded neurons (see above).

#### Recording and characterization of voltage-gated currents

The expression in CSF-cNs of voltage-gated sodium (Na^+^; Na_V_), potassium (K^+^; K_V_) and calcium (Ca^2+^; Ca_V_) channels was assessed in voltage-clamp mode and specific ionic conductances isolated by specifically adapting the composition of the aCSF and the intracellular solution (see ***Supplementary Tables 1 and 2***).

Potassium, sodium and Calcium voltage-dependent currents were elicited while recording neurons at a holding potential (V_*h*_) of -80 mV or -60 mV and voltage steps (*V*_*Step*_) protocols applied to selectively activate a given ionic conductance (see *Results and Figures* for more details). Briefly, sodium voltage-gated channels were activated from a V_*h*_ of -80 mV and *V*_*Step*_ applied from -80 to +60 mV in 10 mV increments applied for 100 ms. Potassium voltage-dependent currents were elicited from V_*h*_ -80 mV with a voltage pre-step to -100 mV for 50 ms to ensure full reactivation of fast inactivating currents (*i*.*e*. I_A_) followed by 100 ms V_*Step*_ ranging from -100 to +60 mV in 10 mV increments before stepping back to -80 mV. To activate I_Ca_, neurons were recorded either from V_*h*_ set at -60 mV with voltage steps applied for 100 ms (*V*_*Step*_, -40 mV to +30 mV, 10 mV increments) or a voltage-ramp protocol from a V_*h*_ of -60 mV and -80 mV (V_*h*_ -60 mV: ramp from -60 to +30 mV over 50 ms and 1.8 mV/ms; V_*h*_ -80 mV: 50 ms pre-step to -120 mV followed by a ramp from -120 to + 30 mV over 100 ms and 1.5 mV/ms). At the end of each recording, the analyzed conductance was blocked using a specific blocker (see ***Supplementary Tables 1 and 2***) to confirm its nature.

#### Recording of ligand-gated-mediated currents

To assess for the expression of GABAergic (GABA_A_-Rs), glycinergic (Gly-Rs), glutamatergic (Glu-Rs) and nicotinic cholinergic (nACh-Rs) ionotropic synaptic receptors, CSF-cNs (lumbar to cervical) were recorded at a V_*h*_ of -80 mV and selective agonists applied by pressure using a patch pipette (∼1 µM tip diameter), positioned at ∼50 µm from the recorded cells and connected to pressure application system (Toohey Company, Picospritzer or WPI). We tested application of γ-amino butyric acid (GABA, 1 mM for 30 ms), glycine (Gly, 1 mM for 100 ms), acetylcholine (ACh, 4 mM for 500 ms), Glutamate (Glu, 100 µM for 100-500 ms). The recordings were performed in the presence of a cocktail of selective antagonists to selectively target one specific ionotropic receptor (see text for more details). To identify the glutamatergic ionotropic receptor subtypes potentially expressed in CSF-cNs, we tested application of α-amino-3-hydroxy-5-methyl-4-isoxazolepropionic acid (AMPA; 100 µM for 100-500 ms), Kainate and N-methyl-D-aspartate acid (NMDA; 100 µM for 100-500 ms). We also tested expression of ionotropic purinergic receptors using pressure application of ATP-γ-S (1 mM for 100-500 ms) but failed to elicit any current. To confirm the nature of the tested receptors, we applied at the end of the recordings the agonist in the presence of the selective antagonist (see ***Supplementary Tables 1 and 2***).

#### Modulation of voltage-gated calcium currents by G-Protein Coupled Receptors

To assess whether postsynaptic voltage-gated Ca^2+^channels are modulated by G-Protein Coupled Receptors (GPCRs), CSF-cNs were recorded in voltage-clamp mode using a specific intracellular solution (see ***Table 1***). Slices were perfused with oxygenated aCSF supplemented with 0.5 µM TTX, and 20 mM TEA-Cl to selectively isolate Ca^2+^currents. Ca^2+^currents were elicited from a V_*h*_ of -80 mV at the peak amplitude with a 100 ms voltage step to +10 mV preceded by a prepulse to -120 mV for 50 ms. Currents were recorded every 20 s over a 5 min period before (Control, CTR), during (Drug) and after (Wash) pressure application of a given GPCR agonist: baclofen (Bcl: 100 µM, 40 s) for GABA_B_ receptors (GABA_B_-Rs), oxotremorine-M (OxoM; 100 µM, 50 s) or acetylcholine (ACh; 1 mM, 40 s) for muscarinic cholinergic receptors (mACh-Rs). We also tested I_Ca_ modulation by metabotropic glutamate receptors (mGlu-Rs) using glutamate pressure application (Glu; 100 µM or 1 mM, 50 s) but failed to elicit any modulation of I_Ca_.

### Reagents

Unless otherwise stated, all chemicals and drugs were purchased from Sigma-Aldrich. Selective blockers or antagonists were applied to block/inhibit channels and receptors as follows: Na_V_ with tetrodotoxin (TTX, T-550, Alomone labs); K_v_ with tetraethylammonium (TEA) and 4-aminopyridine (4-AP); Ca_V_ with cadmium (Cd^2+^, Ca_V_ blocker); GABA_A_ receptors with gabazine (Gbz, SR 95531; Ascent Scientific) and Picrotoxine; glycine with strychnine (Stry, Ascent Scientific); glutamate receptors with 6,7-dinitroquinoxaline-2,3-dione disodium salt (DNQX, Tocris Bioscience); nACh-Rs with D-tubocurarine (D-Tubo) Tocris Bioscience); GABA_B_-Rs with CGP54626A (CGP, Novartis Pharma AG); mACh-Rs with atropine (Atr). For the agonists: oxotremorine-M (Oxo-M, Tocris Bioscience), ATP-γ-S (Abcam Biochemicals) Baclofen (Bcl, Tocris), acetylcholine (ACh, Tocris Bioscience), α-amino-3-hydroxy-5-methyl-4-isoxazolepropionic acid (AMPA, Tocris Bioscience), Kainate and N-methyl-D-aspartate acid (NMDA, Tocris Bioscience), Kainate (Tocris Bioscience) and N-methyl-d-aspartate acid (NMDA, Tocris Bioscience), γ-aminobutyric acid (GABA, Tocris Bioscience), glycine (Gly, Sigma-Aldrich). Experiments were also performed using pressure application of aCSF in the absence of drugs to test for the artefacts or mechanicals responses. However, no changes in the holding current were observed in any of those control experiments confirming that effects were drug or test solution specific. More details about the experimental procedures and solution used are available in ***Supplementary Tables 1 and 2***.

### Processing, analysis and statistics

#### Cell counting in Light sheet microscopy images

Segmentation and cell counting in light sheet microscopy images was performed with FIJI (see ***Supplementary Figure 4*** for graphical explanation and workflow). Briefly, a rolling ball background subtraction of exactly the size of CSF-cNs soma diameter was used (∼10 pixels). Afterwards, a difference of Gaussian filter in 2D was used (Sigma1=1, Sigma2=2) using the CLIJ2^53^ version of “Difference of Gaussian Filter”. Then, the “H-Maxima Local Maximum detection” plugin (from the SCF-MPI-CBG plugins) was used using a Threshold=1 and a Minimum grey value distance between two local optima=5. To remove the H-Maxima ascribable to the ventral axon bundles of CSF-cNs, a 3D signal of the axon bundles was created over the original images using the CLIJ version of the 3D object counter^54^ and these objects loaded into the 3D manager from the 3D ImageJ suite^55^ in order to remove the maxima found within the axon bundles. Afterwards, the remaining particles were filtered by size (<100 voxels) and sphericity in pixel units (>0.5) using the 3D manager, since large and non-spherical particles were found to correspond to very low fluorescence signals. The remaining objects were then plotted on an image stack of the same dimensions than the source one and labeled (as 16 bits images) using “3D simple segmentation” of the 3D ImageJ suite. To visually check for correspondence between the labels and CSF-cNs somas the command “Reduce labels to centroids” from CLIJ2 (see ***Supplementary Figure***) was used. To validate this cell counting strategy, independent acquisitions were made with a 12x objective over a region already imaged with the 4x objective, cells were counted likewise and the total number of cells in this region compared between the two acquisition modes (see ***Supplementary Figure 1***).

#### Cell counting and distribution of CSF-cNs in confocal images

Five 3-6 weeks mice were used for CSF-cNs quantification in confocal slices. 10 fields of view (FOV) were acquired per animal and SC segment (total 30 FOV/animal). To count CSF-cNs in confocal coronal slices, z-stacks of 10-20 images were acquired. Counting was aided by custom FIJI macros available upon reasonable request. In brief, the user was prompted to click in the center of CSF-cNs on a Maximal Projection image while being able to navigate in the z-stack in order to accurately resolved closely packed cells. Once all the cells were marked, the user was asked to define manually the dorsal and ventral regions on the CC (ventral fiber bundles served as bona fide indicators of SC orientation). Then, the number of cells in each quadrant and their percentages were automatically calculated, and the cell densities/10 μm of SC were expressed as (Total number of CSF-cNs in the field of view x 10)/(Number of optical slices x Voxel depth). Afterwards the DAPI signal was automatically thresholded to create the contour of the CC, and the Euclidean distance (in microns) of each cell to this contour was calculated using the “Exact Euclidean Distance Transform (3D)” function in FIJI^56^ along with the pixel size of the image.

OriginPro was used for statistical analyses and generating plots. Normality was tested for all conditions. Since at least one group was always found to not meet the normality assumption in every possible comparison, Kruskal-Wallis ANOVA followed by paired post-hoc *Dunn* test was used for cells densities and distributions. Results were considered significant when *p<0.05, **p<0.01. For comparing distances of CSF-cNs to the CC at different SC levels, Kolmogorov–Smirnov test was used. Since K-S test is extremely powerful over large data sets, results were considered significant when *p<0.001.

#### Electrophysiology

Passive properties were determined, in voltage-clamp mode at -70 mV, from the current responses to a -10 mV hyperpolarization step (*V*_*Step*_). Membrane resistance (*R*_*m*_) was calculated from the amplitude of the sustained current at the end of the voltage step (*R*_*m*_*=V*_*Step*_*/I*_*m*_). Membrane capacitance (*C*_*m*_) was estimated as the ratio between the cell decay time constant (*τ*_m_), obtained from the exponential fit of the current decay, divided by *R*_*s*_ (*C*_*m*_ *∼ τ*_m_*/R*_*s*_). Resting membrane potential was determined in current-clamp mode at I=0 just after the whole-cell configuration was achieved. AP discharge pattern was analyzed from trains of APs elicited by the injection DC current pulses using the ‘threshold detection’ routine from Clampfit 9-11 with a threshold set at 0 mV. Voltage and current responses to agonists were analyzed using the Clampfit 9-11 suite (Molecular Devices Inc.) and Microsoft Excel 2018-2024.

PKD2L1 unitary current analysis was conducted using the WinEDR v4.0.3 software (Strathclyde University) where all points histograms for the close (baseline) and the open (PKD2L1 unitary currents) states were generated on 500 bins over 20-30 s recording periods to determine the respective mean current. Histogram were fitted with gaussian curves defined by the following equation:

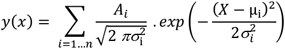

Where A_*i*_ corresponds to the area under the curve, *X* the current amplitude, µ_*i*_ the mean current and σ_*i*_ to the standard deviation. PKD2L1 unitary current amplitudes were calculated as the difference between the mean current values (µ_*i*_) of the open and close states. The open probability (NPo) was computed from the area under the curve of the open state histogram. Data were analyzed in control, during pH solution application and following wash and compared. For ASICs, the peak amplitude was measure after subtracting the baseline current. Modulation of the spiking properties following exposure to pH solution, were analyzed from the RMP and the instantaneous AP frequency determined using the Clampfit (Molecular Device, Inc.) threshold detection routine.

For Na_V_ and K_V_, the current-voltage relationship for a given conductance was constructed for each cell by plotting the Na^+^ and K ^+^ current peak amplitude recorded for a given V_*Step*_, and the mean IV curve generated either by pooling the data for all cells recorded from the 3 segments. Ca_V_ mediated currents were elicited from V_*h*_ -60 mV or -80 mV with the voltage-ramp against time protocol. The Ca_V_ IV-curve was subsequently obtained by converting the data as current against voltage. For each level, to compare recordings elicited from V_*h*_ -80 and -60 mV and distinguish HVA and LVA channel activation, the holding current was subtracted from individual converted IV-curves, and average curves calculated and subsequently normalized to the peak (current amplitude are presents as arbitrary unit, AU). In the current traces represented to illustrate specific voltage-dependent channel activation against voltage, remaining capacity transients were digitally subtracted.

All data are expressed as mean±SD or SEM, as specified, and N and n correspond to the number of animal and data sample, respectively. They are represented with boxplots and whisker using the Tukey’s method as well as single points to illustrate data distribution. In boxplots with whisker, for each data set, the median (represented as thick line) and the 25th to the 75th percentiles (lower and upper limits of each bar, respectively) are calculated. Next, the interquartile distance (IQR) is determined as the difference between the 25th and 75th percentiles, and the whisker limits or “inner fences” calculated as the 75th percentile plus 1.5 times IQR and the 25th percentile minus 1.5 times IQR. All data with either higher or lower than the inner fences are represented as individual data points and considered as outliers (circles on the plots).

Statistical analyses were carried out using the R Studio 3.3.1 statistical software (R Studio Team, 2015) in the R environment. Since, data did not meet the normality assumption (Shapiro normality test), we used non-parametric statistical tests. For results where data are only compared for one variable, we used *Kruskal-Wallis* one-way ANOVA statistical analysis and provide 𝒳^2^, degree of freedom (number of parameters-1, df), R^2^ and p(𝒳^2^) values, for the analyzed factor. To compare data within a SC segment in different condition (control, CTR *vs*. application of agonists or blockers) as well as between segments along the rostro-caudal axis, the “*non-linear mixed effect*” (nlme version 3.1–128 package^57^) was used. The distinction between CSF-cNs response in the different experimental conditions is considered a *Condition* variable, while the distinction between the different neurons along the CC rostro-caudal axis represents the *Region* variable. On one hand, in one neuron for a given Region, its response in one Condition (control *vs*. drugs application or wash) is assumed dependent of the other recording Condition, and the variable represents a dependent factor. On the other hand, the same variables represent independent factors when considered as a function of the Region (cervical, thoracic and lumbar). Therefore, the statistical analyses required using a mixed effect model considering in a hierarchical way both dependent (*Condition*) and independent variables (*Region*) as well as the interaction between the variables. The null hypothesis was set as the absence of difference between the data among each group. The nlme model was tested by using an analysis of the variance (ANOVA) and we provide the F-statistic (F), the degree of freedom (df) and the p(F) values for the given fixed factor tested (*Condition, Region*). Wherever an interaction between the *Region* and *Condition* factors was statistically significant, we reported the p-value. We further conducted a post-hoc *Tukey (EMM) post-hoc test* to test for Contrasts, pairwise comparisons and interaction as a function of the *Region* and the *Condition*. Statistical differences were considered as significant for * p < 0.05.

## ADDITIONAL INFORMATION

### Data availability statement

The data of this manuscript are available from the corresponding author upon reasonable request.

## Authors contributions

All experiments were performed in laboratories at the ‘Institut des Neurosciences de la Timone (INT)’ of Aix-Marseille University and CNRS, UMR 7289. Conception and design of the work: CE, BE, JN, RFJ, SR & WN. Acquisition, analysis, and interpretation of data for the work: CE, BE, JN, RFJ, SR, MC & WN. Drafting the work or revisiting it critically for important intellectual content: CE, BE, JN, RFJ, SR, MC, JT & WN. All authors have read and approved the final version of this manuscript and agree to be accountable for all aspects of the work in ensuring that questions related to the accuracy or integrity of any part of the work are appropriately investigated and resolved. All persons designated as authors qualify for authorship, and all those who qualify for authorship are listed.

## Acknowledgments

We acknowledge the ‘Institut de Neurosciences de la Timone (INT)’ technical facilities for their support in the study (*Neuro-Bio-Tools*: Molecular Biology (NeuroVir)/Histology (ConnectoVir) and *Photonic Neuroimaging* (INPHIM, light-sheet & confocal microscopy) as well as the *Mediterranean Primate Research Centre* (MPRC; UAR 2018), Animal facility).

---

**Supplementary Figure 1.**
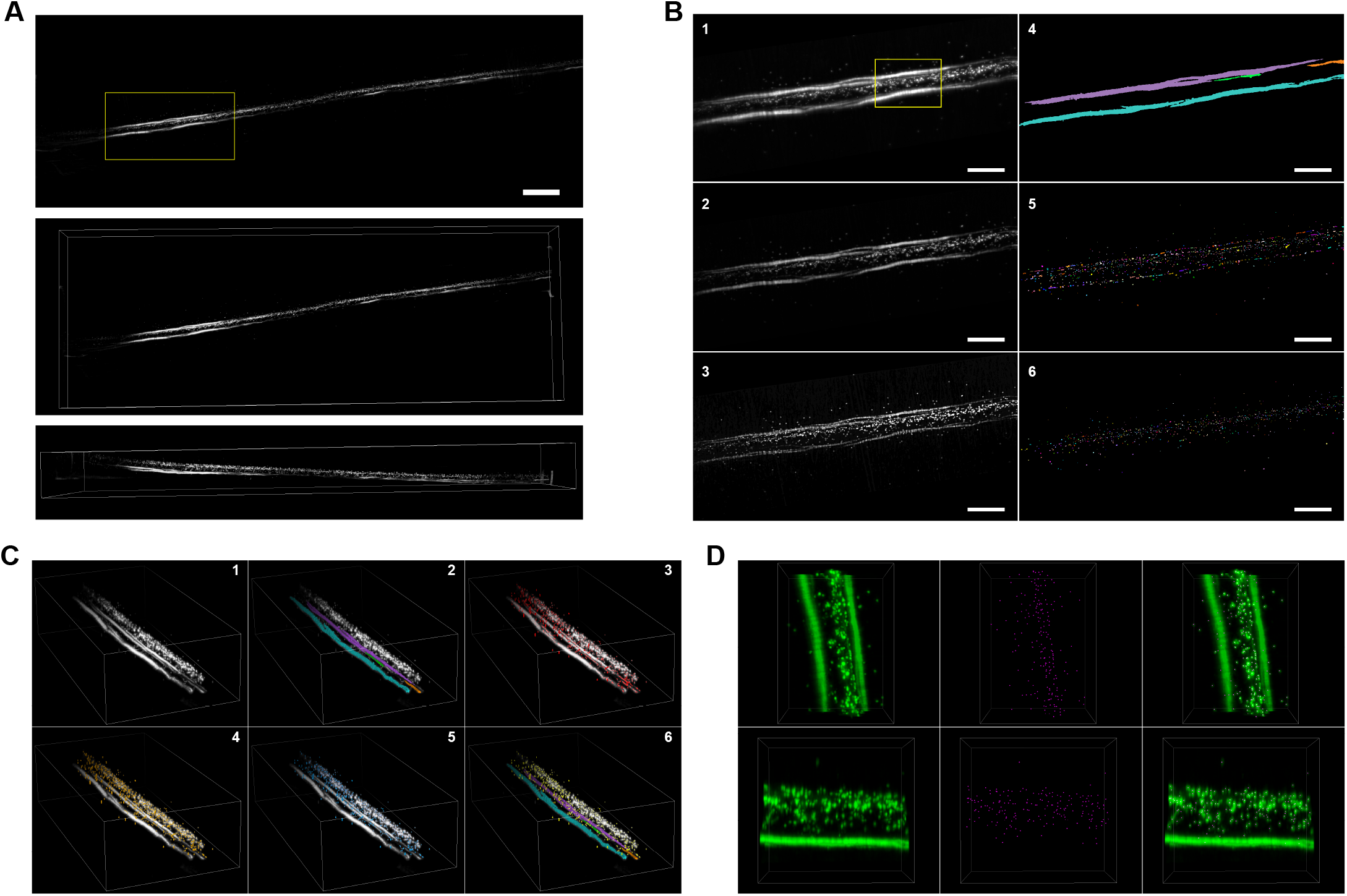
Comparison of the cell segmentation workflow and estimation of cell number at different light-sheet resolutions. **A)** Full view corresponding to **Fig. 1A**, showing regions acquired at different light sheet resolutions (1x, full view; 4x, large box; 12x small box). **B)** Detailed view of the thoracic region boxed in **Supplementary Figure 1A**, showing the estimation for cell numbers at 4x (yellow) and 12x (green) in equivalent light-sheet microscopy stacks. Boxes sizes are in μm along the Rostro-caudal (RC) x Latero-Median (LM) x Dorso-Ventral (DV): 4x image (yellow) 3076.13 × 1784.25 × 818; 12x image (green) 692.90 × 703.95 × 211.25. **C)** Maximal projection of the region boxed in **Supplementary Figure 1B** in fluorescence (Top) and particle analysis of this same region (Bottom). Scale bar=50 μm. **D)** 3D views of the same region of the maximal projection depicted in **C** from lateral (**Left**), Top (**Middle**), and front views (**Right**). Fluorescence images are depicted in green (**Top**) and the result of the particle analysis workflow is merged in magenta (**Bottom**). CC: Central canal. Scale: see 1B.

**Supplementary Figure 2.**
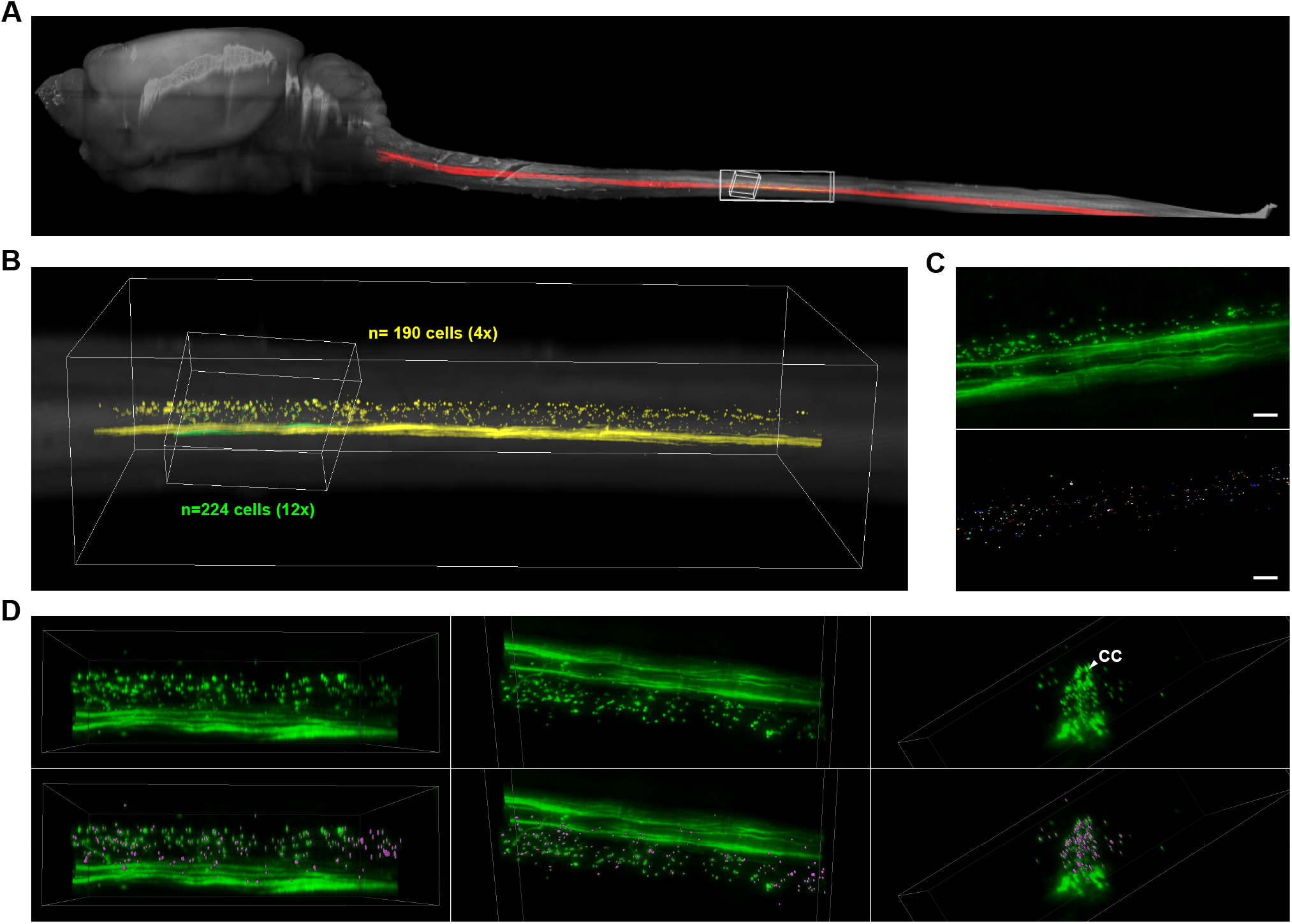
Size comparison of CSF-cNs to motor neurons using different microscopy techniques. **A)** Confocal images of a large field of view showing CSF-cNs expressing EGFP (green) in a ChAT-tdTomato mouse (ChAT^+^, in magenta and DAPI in blue). Expression of EGFP is triggered by intra cerebroventricular (ICV) injections of AAV1-hSyn-EGFP particles as previously described. Scale bar=50 μm. **B)** Detailed views of the field of view shown in A corresponding to the boxes depicted in the merge panel of A (Top). Scale bar=20 μm. **Bottom, Left**. 3D renderings of the neurons depicted in Top panels. **Bottom, Right**. silhouette of one CSF-cN (green) and one motor neuron (MN, magenta) for size comparison. **C)** Light-sheet microscopy images obtained at the highest possible magnification of the light-sheet microscope in ChAT-tdTomato animal showing MNs and other ChAT^+^ neurons (magenta), and in a PKD-tdTomato animal showing CSF-cNs (green). Note that even at the highest magnification, the resolution of the light-sheet microscope combined to the DISCO clearing technique remains poor to resolve individual CSF-cN somatas. Boxes sizes are (RC × LM × DV in μm); for ChAT^+^ neurons: 555 × 752 × 468; for CSF-cNs: 565 × 285 × 266 (in µm).

**Supplementary Figure 3.**
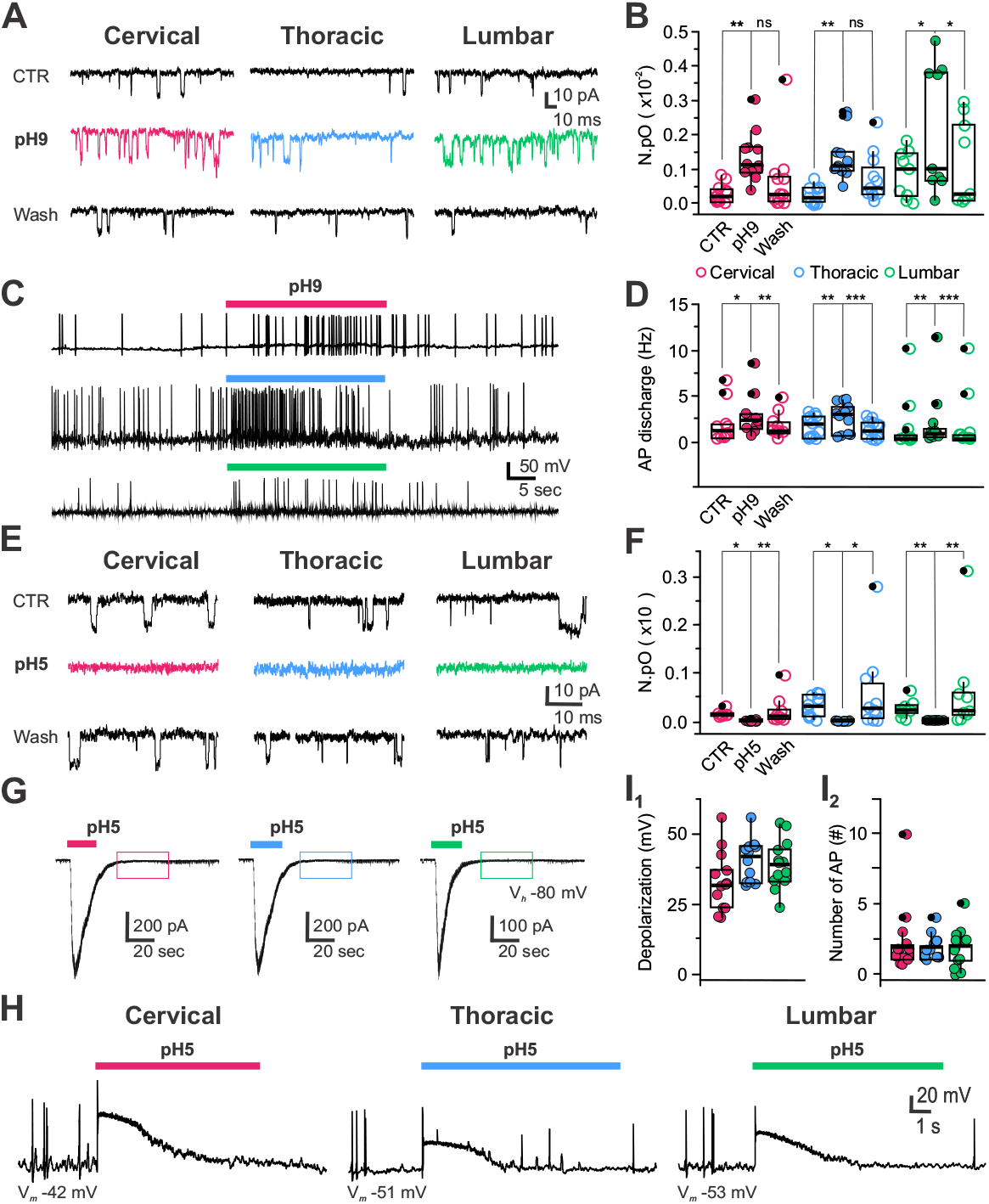
CSF-cNs have chemosensory properties through modulation of PKD2L1 channels and ASICs. **A)** Representative recordings in voltage-clamp mode at V_*h*_ -80 mV of PKD2L1 channel activity in CSF-cNs from the cervical, thoracic and lumbar regions (***Left to Right***) in control (Top), upon exposure to extracellular alkalinization (pH 9; colored traces; ***Middle***) and following washout (Wash, ***Bottom***). **B)**Summary plots for the average channel open probability (NPo) in control (CTR), in pH 9 and after Wash for CSF-cNs from the cervical (red), thoracic (blue) and lumbar (green) segments (CTR: 0.027±0.027, 0.025±0.022 and 0.085±0.069; pH 9: 0.0.134±0.071, 0.135±0.068 and 0.215±0.183 and Wash: 0.064±0.100, 0.074±0.070 and 0.115±0.126 for C (N=3, n=12), T (N=6, n=11), and L Segments (N=4, n=9), respectively; Data are multiplied by 100 on the graph for a better visualization; *ANOVA*.*lme*: F=4.52, df=8 and 87, p(F)= 0.0001272 and *Tukey (EMM) post-hoc* test to compare CTR *vs*. pH 9, CTR *vs*. Wash and pH 9 *vs*. Wash within the different regions and between region; * and ** for p<0.05 and <0.01, respectively). **C)** Representative recordings in current-clamp mode at RMP (C: -59 mV; T: -47 mV and L: - 41 mV) of CSF-cN AP firing activity in control, during pH 9 exposure (colored bars) from cervical, thoracic and lumbar segments (***Top to Bottom***). **D)** Summary plots for the average instantaneous AP frequency in control (CTR), in pH 9 and after Wash for CSF-cNs from the cervical (red), thoracic (blue) and lumbar (green) segments (CTR: 1.8±2.0, 1.5±1.2 and 1.4±2.9 Hz; pH 9: 2.8±2.4, 2.6±1.6 and 1.9±34.2 and Wash: 1.7±1.4, 1.2±0.9 and 1.5±3.1 Hz for C (N=4, n=10), T (N=5, n=13), and L Segments (N=3, n=12), respectively; *ANOVA*.*lme*: F=18.52, df=8 and 96, p(F)=2.2.10^−16^ and *Tukey (EMM) post-hoc* test to compare CTR *vs*. pH 9, CTR *vs*. Wash and pH 9 *vs*. Wash within the different regions and between region; *, **, *** for p<0.05, <0.01 and <0.001, respectively). **E)** Representative recordings in voltage-clamp at V_*h*_ -80 mV of PKD2L1 channel activity in CSF-cNs from the cervical, thoracic and lumbar regions (***Left to Right***) in control (***Top***), upon exposure to extracellular acidification (pH 5; colored traces; ***Middle***) and following washout (Wash, ***Bottom***). **F)** Summary plots for the average channel open probability (NPo) in control (CTR), in pH 5 and after Wash for CSF-cNs from the cervical (red), thoracic (blue) and lumbar (green) segments (CTR: 0.015±0.006, 0.031±0.022 and 0.025±0.018; pH 5: 0.0.1±0.002, 0.001±0.001 and 0.000±0.000 and Wash: 0.022±0.030, 0.058±0.085 and 0.062±0.097 for C (N=3, n=9), T (N=4, n=10), and L Segments (N=3, n=9), respectively; Data are multiplied by 100 on the graph for a better visualization; *ANOVA*.*lme*: F=2.499, df=8 and 75, p(F)= 0.01835 and *Tukey (EMM) post-hoc* test to compare CTR *vs*. pH 5, CTR *vs*. Wash and CTR *vs*. Wash within the different regions and between region; * and ** for p<0.05 and <0.01, respectively). **G)** Representative current traces elicited in CSF-cNs recorded in voltage-clamp mode at V_*h*_ -80 mV in the cervical, thoracic and lumbar segments (***Left to Right***) upon pressure application of an acidic solution (pH 5, colored bars). The colored boxes during the persistent phases of the current illustrates the absence of PKD2L1 activity (see also ***Supplementary Figure 3E***). **H)** Representative voltage traces recorded in current-clamp mode at RMP (in CSF-cNs from the cervical, thoracic and lumbar segments (***Left to Right***) upon pressure application of an acidic solution (pH 5, colored bars). **I)** Summary plots for the average amplitude of the depolarization induced (**I**_**1**_) and the number of AP triggered (**I**_**2**_) following exposure to acidic pH in CSF-cNs from the cervical (red), thoracic (blue) and lumbar (green) segments. (**I**_**1**_) Depolarization: +33±10 mV, +40±8 mV and +39±9 mV for cervical (N=5, n=13), thoracic (N=3, n=12) and lumbar segments (N=3 n=13) (*Kruskal-Wallis rank sum test*: 𝒳^2^=4.9908, df=2, p(𝒳^2^)= 0.08246 and a posthoc *Pairwise comparisons using Wilcoxon rank sum test* with continuity correction: C *vs*. T, p=0.12; C *vs*. L, p=0.12 and T *vs*. L, p=0.81). (I_2_) Number of PA: 2.4±2.5, 1.8±0.9 mV and +1.8±1.4 mV for cervical (N=5 n=13), thoracic (N=3, n=12) and lumbar segments (N=3 n=13); *Kruskal-Wallis rank sum test*: 𝒳^2^=0.074495, df=2, p(𝒳^2^)= 0.9634 and a posthoc *Pairwise comparisons using Wilcoxon rank sum test* with continuity correction: C *vs*. T; C *vs*. L and T *vs*. L, p=0.98).

**Supplementary Figure 4.**
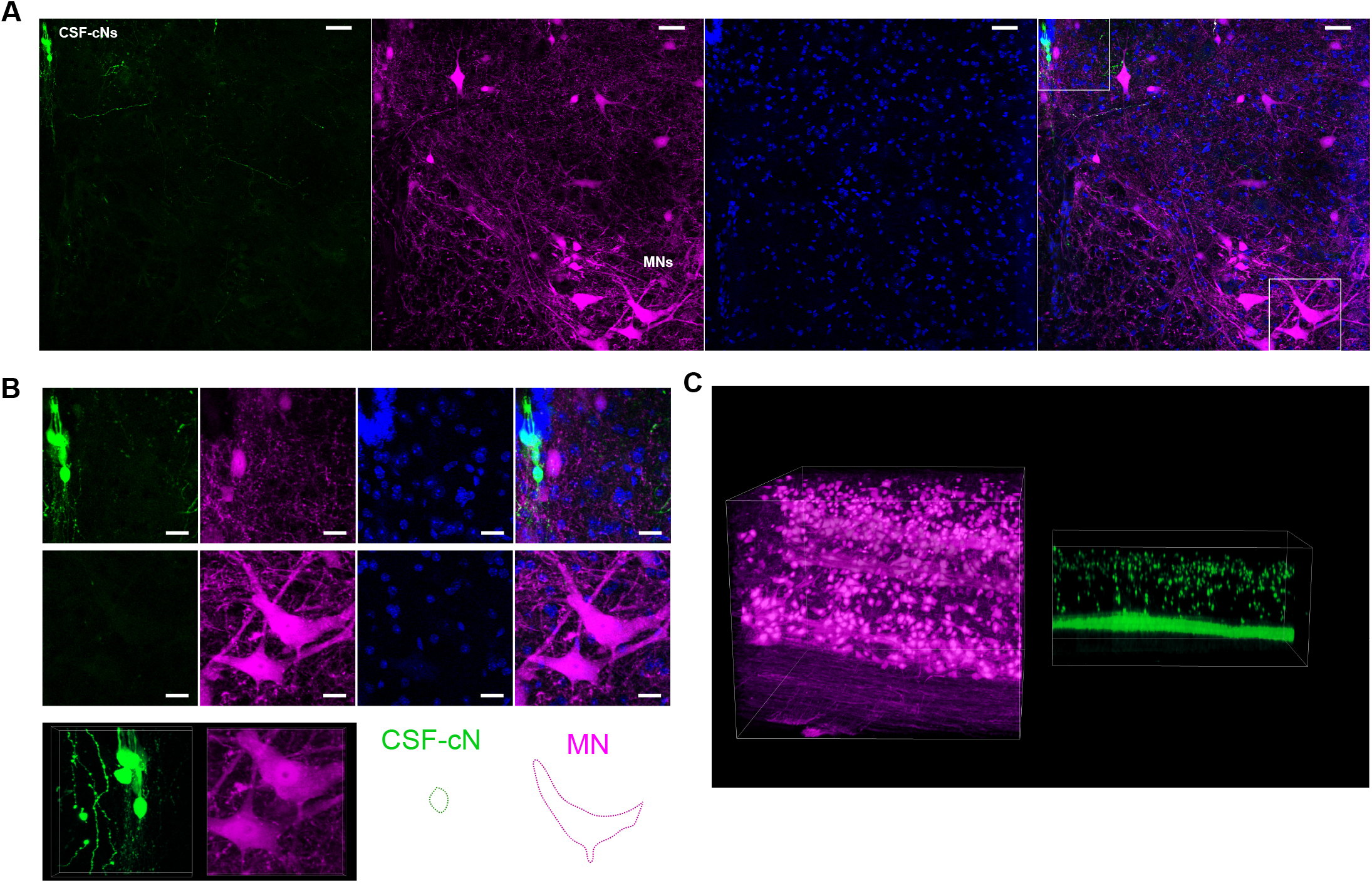
Workflow for cell segmentation and estimation of cell number. **A)*Top***. Maxima*l* projection of a segment corresponding to the cervical SC of a PKD-tdTomato mouse. The inset box corresponds to the region depicted in ***Fig. 1B*** and ***Supplementary Figure 1B. Middle***, dorsal view of the 3D rendering for the region depicted in the Top panel. ***Bottom***. lateral view of the 3D rendering of the region depicted in the Top panel. Scale bar=500 μm. Dimensions of the box are (RC × LM × DV in μm) 7484.75 × 2632.50 × 841.75. **B)** Graphic workflow for image segmentation of the inset in ***Supplementary Figure 1A*** (see methods). 1) Maximal projection of the inset shown in ***Supplementary Figure 1A***. 2) After rolling ball background subtraction r=10 pixels. 3) After Difference of Gaussians filter (sigma1=1, sigma2=2). 4) 3D objects corresponding to the ventromedial axon bundles of CSF-cNs. 5) H-Maxima obtained from the image depicted in 3). 6) H-Remaining particles after removing H-Maxima belonging to ventromedial axon bundles and filtering by volume and sphericity. Scale bar=200 μm. **c)** 3D representations of the views depicted in ***Supplementary Figure 1B***. 1) Corresponds to ***Supplementary Figure 1B***, 1. 2) Corresponds to ***Supplementary Figure 1B***, 4. 3) corresponds to ***Supplementary Figure 1B, 5***. 4) is ***Supplementary Figure 1B, 5*** after removing particles located in the ventral axon bundles. 5) After filtering by volume. 6) Merge of the final result of particle quantification and segmentation of the axon bundles. Dimensions of the box are (RC × LM × DV in μm) 1764.75 × 913.25 × 463.00. **D)** Inset of the box depicted in ***Supplementary Fig. 1B*** in dorsal (***Top***) and lateral (***Bottom***) views. ***Left***. Fluorescence image of CSF-cNs. ***Middle***. Centroids corresponding to particle analysis of this specific region. ***Right***. Merge of the centroids from particle analysis to the fluorescence microscopy images.

