## Supplementary figures and images for "Spinal cerebrospinal fluid contacting neurons are a conserved sensory neuronal population along the mouse spinal cord"

### Supplementary Fig 1

#### Crozat, Blasco et al., Figure Supplementary 1

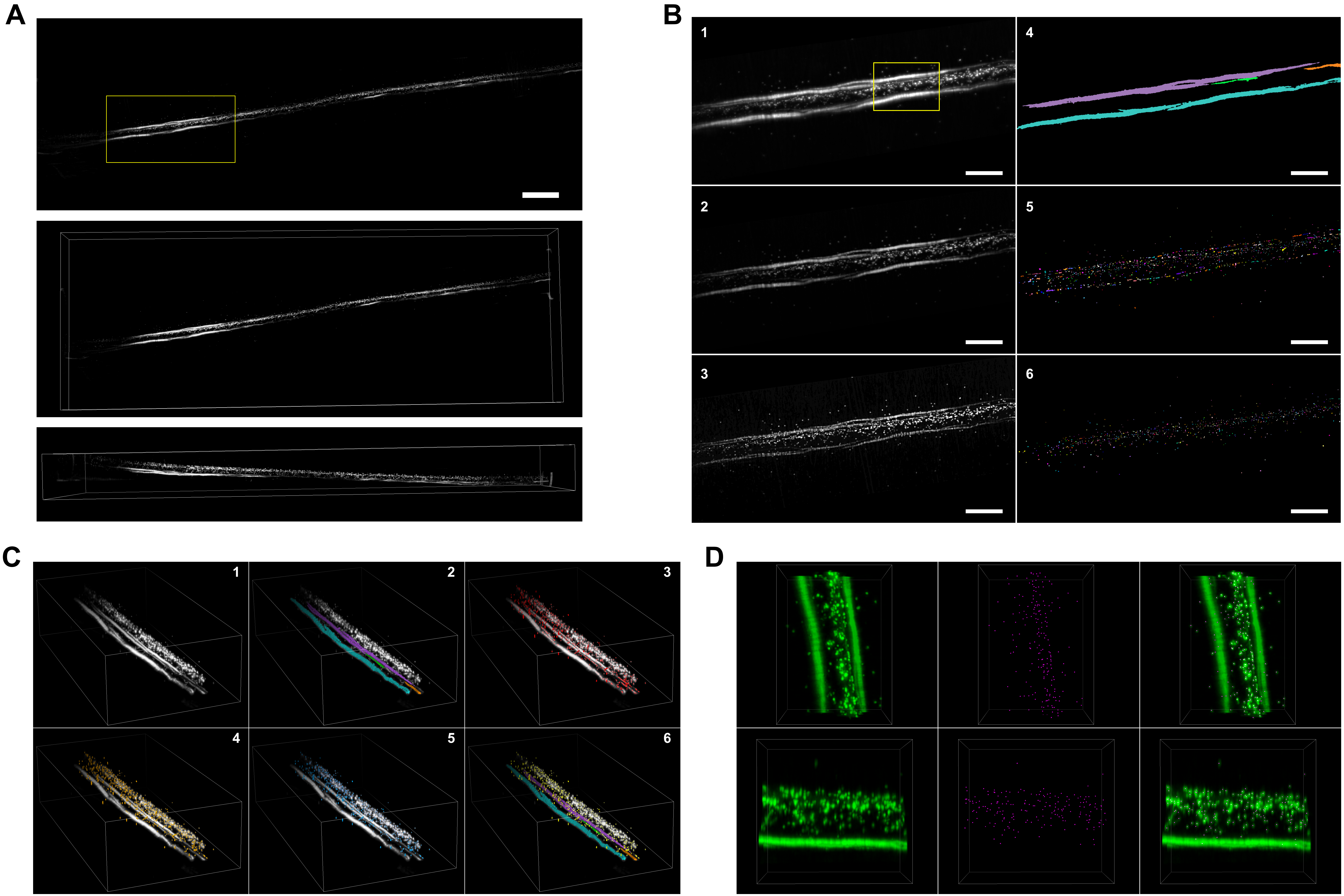

### Supplementary Fig 2

#### Crozat, Blasco et al., Figure Supplementary 2

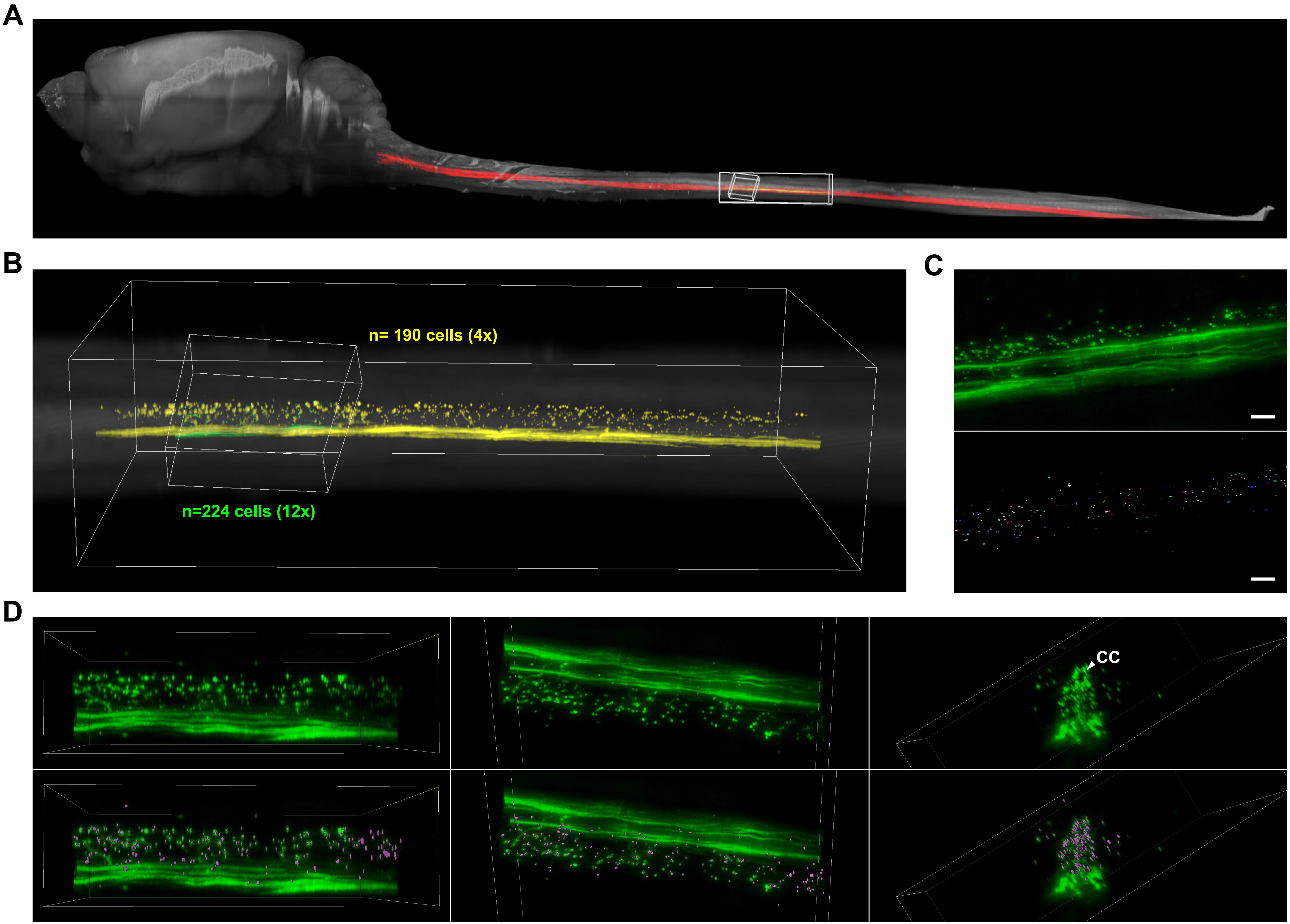

### Supplementary Fig 3

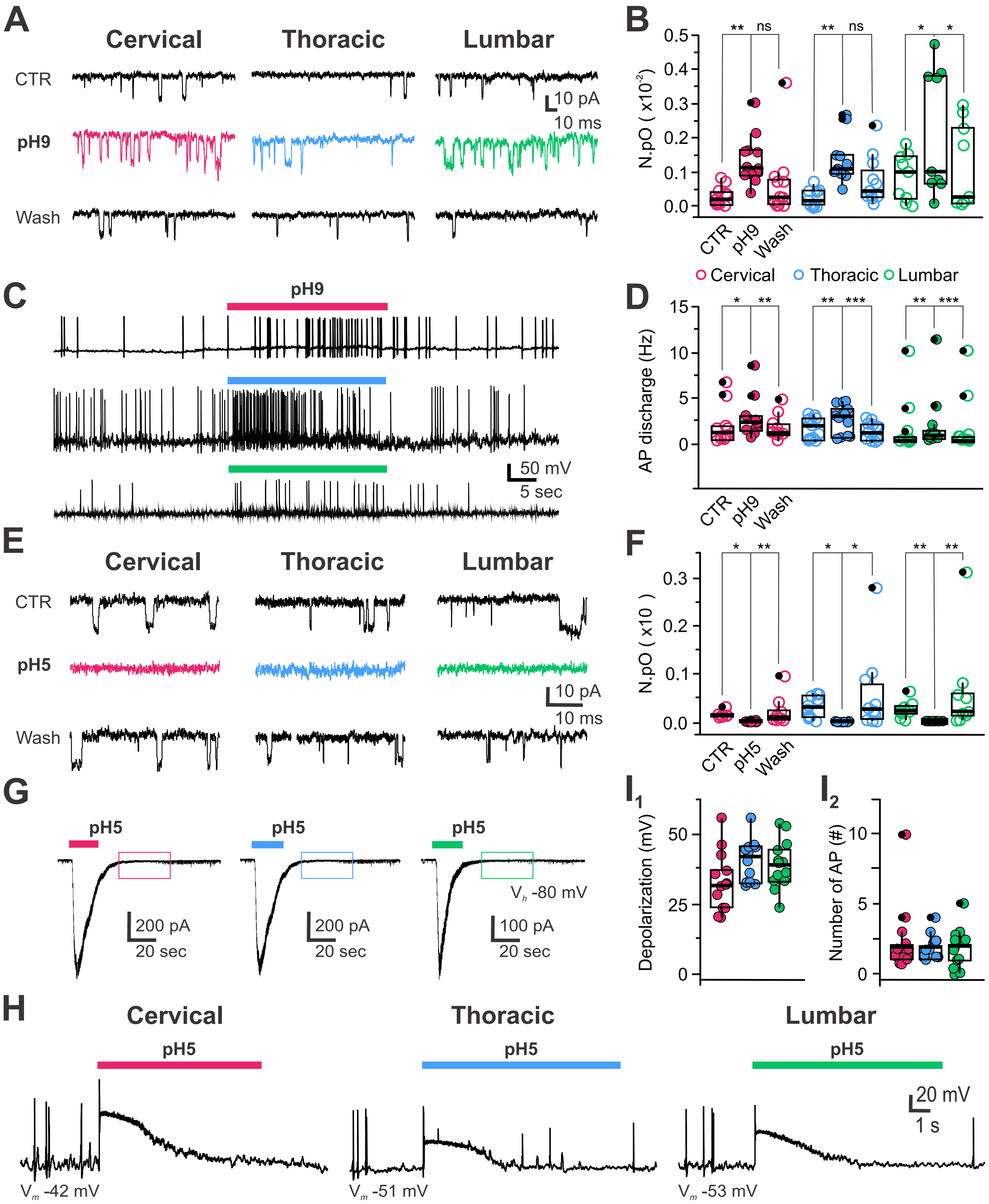

### Supplementary Fig 4

Crozat, Blasco et al., Figure Supplementary 3

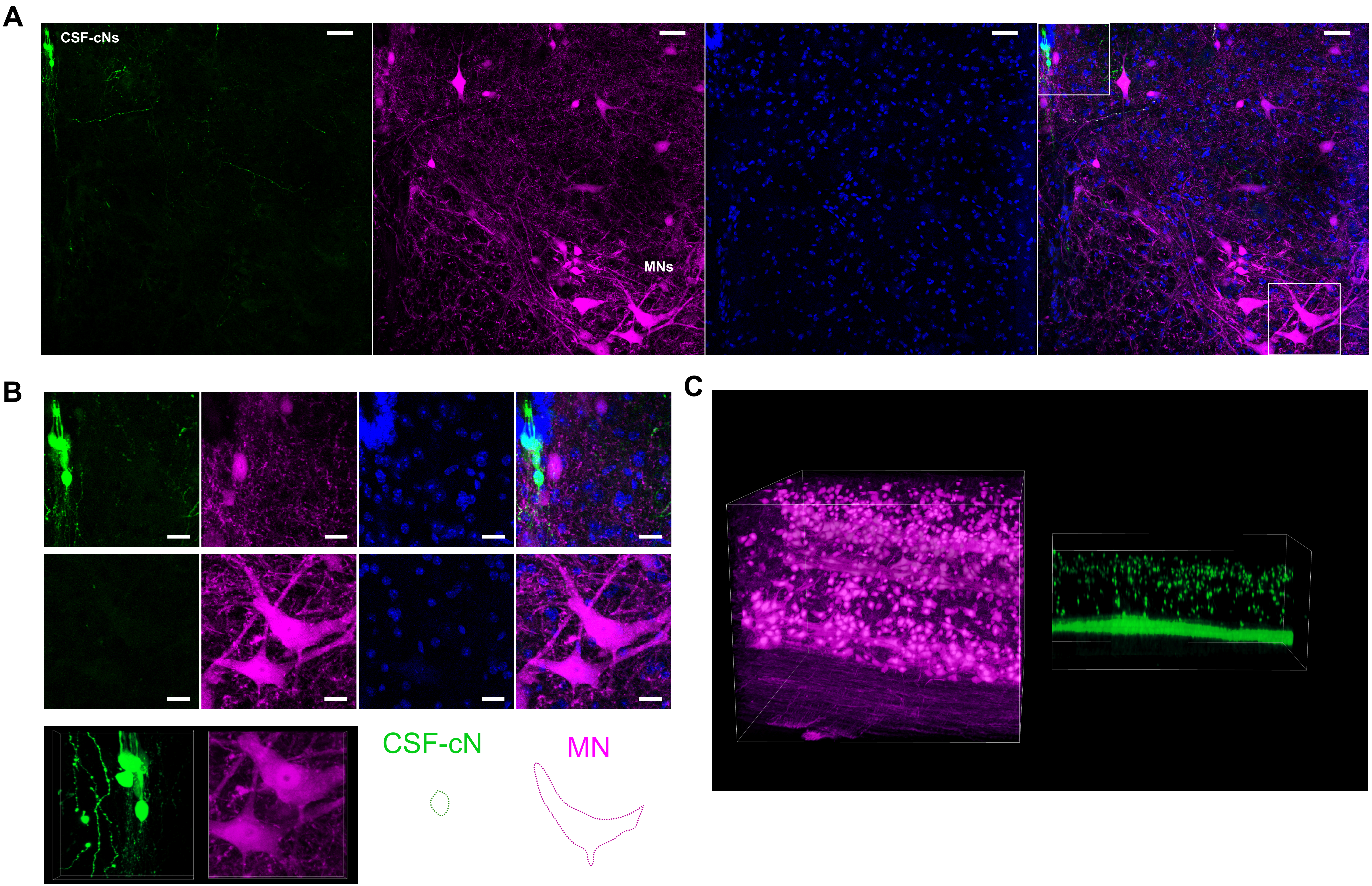
