## Supplementary Table 1 for "Spinal cerebrospinal fluid contacting neurons are a conserved sensory neuronal population along the mouse spinal cord"

### Table S1 - Composition of the intracellular solutions used for the sets of recording

| **Major Ion** | **KGlu (A)** | **CsAc-TEA (B)** | **KCl (C)** | **CsAc (D)** |
| --- | --- | --- | --- | --- |
| **Composition (mM)** | | | | |
| **KCl** | 3 | - | 130 | - |
| **K-gluconate** | 120 | - | - | - |
| **NaCl** | 5 | 5 | 10 | 5 |
| **CsCl** | - | 3 | - | 3 |
| **CsAc** | - | 100 | - | 120 |
| **TEA-Cl** | - | 20 | - | - |
| **MgCl_2_** | 1 | 1 | 2 | 1 |
| **CaCl_2_** | 1 | 0.5 | 1 | 1 |
| **Free [Ca^2+^]*_i_*** | 27 nM | 12 nM | 13 nM | 12 nM |
| **EGTA** | 2 | - | 5 | 10 |
| **BAPTA** | - | 10 | - | - |
| **HEPES** | 10 | | | |
| **Phosphocreatine** | 10 | | | |
| **Mg-ATP** | 4 | | | |
| **Na_2-_GTP** | 0.2 | | | |
| **pH** | 7.35 | 7.35 | 7,35 | 7.34 |
| **pH adjustment** | KOH 8N | CsOH 50% Wt/Vol | KOH 1M | CsOH 50% Wt/Vol |
| **Osmolarity** | ~295 mOsmol.Kg^-1^ | | | |
| **Equilibrium potentials (mV) at 20-25°C** | | | | |
| **E_Na_** | +46 | +46 | +69 | +44 |
| **E_K_** | -94 | - | -97 | / |
| **E_Cl_** | -65 | -64 | +5 | -59 |
| **E_Ca_** | +62 | +65 | +159 | +107 |
