## Supplementary Table 2 for "Spinal cerebrospinal fluid contacting neurons are a conserved sensory neuronal population along the mouse spinal cord"

### Supplementary Table 2 - Composition of the extracellular solutions used to selectively isolate ionic conductances

| **Experiments** | **Solution** | **Solution Label** | **Addition** | **Blockers** |
| --- | --- | --- | --- | --- |
| ***Characterization of intrinsic and AP discharge properties*** | | | | |
| **Passive properties** | Intracellular | A-D  (A for CC) |  |  |
|  | Extracellular | aCSF |  |  |
| ***Characterization of voltage-dependent channels*** | | | | |
| **Na^+^ Current** | Intracellular | B |  |  |
|  | Extracellular | Na_V_ | +TEA (20mM)  without CaCl_2_  replaced by 4mM MgCl_2_ | TTX (0,5 µM) puff |
| **K^+^ Current** | Intracellular | A |  |  |
|  | Extracellular | K_V_ | +TTX (0,5 µM)  without CaCl_2_  replaced by 4mM MgCl_2_ | TEA (10 mM) bath  +4-AP (4 mM) puff |
| **Ca^2+^ Current** | Intracellular | B |  |  |
|  | Extracellular | Ca_V_ | +TTX (0,5 µM)  +TEA (20mM) | Cadmium (200 µM) bath and/or puff |
| ***Characterization of ionotropic ligandd-gated receptors*** | | | | |
| **GABA (1mM)** | Intracellular | C |  |  |
|  | Extracellular | aCSF | +TTX (0.5 µM)  +Stry (1 µM)  +D-tubocurarine  +DNQX (20 µM) | Gbz (10 µM)  +Picrotoxine (100 µM) |
| **Glycine (1mM)** | Intracellular | C |  |  |
|  | Extracellular | aCSF | +TTX (0.5 µM)  +Gbz (10 µM)  +D-tubocurarine  +DNQX (20 µM) | Stry (1µM) |
| **Glutalmate AMPA ; NMDA ; Kainate (100µM)** | Intracellular | C (VC) / A (CC) |  |  |
|  | Extracellular | aCSF | ***Voltage-Clamp :***  +TTX (0.5 µM)  +Gbz (10µM)  +Picrotoxine (100 µM)  +Stry (1 µM)  +D-tubocurarine (100µM) | DNQX (400µM) |
|  |  |  | ***Current-Clamp :***  +Gbz (10µM)  +Picrotoxine (100 µM)  +Stry (1 µM)  +D-tubocurarine (100µM) |  |

| **ACh**  **(4 mM)** | Intracellular | D (VC) / A (CC) |  |  |
| --- | --- | --- | --- | --- |
|  | Extracellular | aCSF | ***Voltage-Clamp :***  +Atropine (10 µM)  +TTX (0.5 µM)  +Gbz (10µM)  +Picrotoxine (100 µM +DNQX (400µM) | D-tubocurarine (100µM) |
|  |  |  | ***Current-Clamp :***  +Atropine (10 µM)  +Gbz (10µM)  +Picrotoxine (100 µM +DNQX (400µM)  +D-tubocurarine (100µM) |  |
| **ATP-γ-S**  **(1 mM)** | Intracellular | C |  |  |
|  | Extracellular | aCSF | +TTX (0.5 µM)  +Gbz (10µM)  +Picrotoxine (100 µM +Stry (1 µM)  +DNQX (400µM) |  |
| ***GPCR modulation of Ca_V_*** | | | | |
| **Baclofen**  **(100 µM, 40s)** | Intracellular | B |  |  |
|  | Extracellular | Ca_V_ Modulation | TTX (0.5 µM)  + TEA (20 mM)  + Strych (1µM  + DNQX (20µM)  + Gbz (10µM) | CGP (2 µM) |
| **OxoM**  **(100µM, 50s)** | Intracellular | B |  |  |
|  | Extracellular | Ca_V_ Modulation | TTX (0.5 µM)  + TEA (20 mM)  + Strych (1µM)  + DNQX (20µM)  + Gbz (10µM) | Atropine (10 µM) |
